# Long-term balancing selection and the genetic load linked to the self-incompatibility locus in *Arabidopsis halleri* and *A. lyrata*

**DOI:** 10.1101/2022.04.12.487987

**Authors:** Audrey Le Veve, Nicolas Burghgraeve, Mathieu Genete, Christelle Lepers-Blassiau, Margarita Takou, Juliette De Meaux, Barbara K. Mable, Eléonore Durand, Xavier Vekemans, Vincent Castric

## Abstract

Balancing selection is a form of natural selection maintaining diversity at the sites it targets and at linked nucleotide sites. Due to selection favouring heterozygosity, it has the potential to facilitate the accumulation of a “sheltered” load of tightly linked recessive deleterious mutations. However, precisely evaluating the extent of these effects has remained challenging. Taking advantage of plant self-incompatibility as one of the best-understood examples of long-term balancing selection, we provide a highly resolved picture of the genomic extent of balancing selection on the sheltered genetic load. We used targeted genome resequencing to reveal polymorphism of the genomic region flanking the self-incompatibility locus in three sample sets in each of the two closely related plant species *Arabidopsis halleri* and *A. lyrata*, and used 100 control regions from throughout the genome to factor out differences in demographic histories and/or sample structure. Nucleotide polymorphism increased strongly around the *S*-locus in all sample sets, but only over a limited genomic region, as it became indistinguishable from the genomic background beyond the first 25-30kb. Genes in this chromosomal interval exhibited no excess of mutations at 0-fold degenerated sites relative to putatively neutral sites, hence revealing no detectable weakening of the efficacy of purifying selection even for these most tightly linked genes. Overall, our results are consistent with the predictions of a narrow genomic influence of linkage to the *S*-locus, and clarify how natural selection in one genomic region affects the evolution of the adjacent genomic regions.

## Introduction

Balancing selection refers to a variety of selective regimes maintaining advantageous genetic diversity within populations (Delph and Kelly 2014). Notable examples include heterozygote advantage, negative frequency-dependent selection, and spatial heterogeneity. In contrast to genetic linkage to genomic sites subject to either positive or negative selection that generally tends to eliminate surrounding genetic variation (Smith and Haigh 1974; Charlesworth et al. 1993), linkage to loci under balancing selection is expected to locally promote the long-term persistence of variation in surrounding sites (Hudson et al. 1987; Charlesworth 2006). Besides the strength of balancing selection and the local rate of recombination, the magnitude of the local diversity increase and its extent along the chromosome critically depend on details of the exact form of balancing selection (Strobeck 1983, Charlesworth et al. 1997, Hudson and Kaplan 1988, Takahata and Satta 1998, Wiuf et al. 2004). An important feature of this form of natural selection is that the extended time over which the balanced allelic lineages are maintained (Takahata and Nei 1990; Vekemans and Slatkin 1994) also means more time for recombination to decouple them from their linked sites, such that the extent of the region affected may end up being quite narrow (Strobeck 1980; Nordborg 1997; Schierup et al. 2001). Genome resequencing studies have revealed that balancing selection can be a potent force throughout the genome (DeGiorgio et al. 2014), but because of inherent technical challenges related to the often high levels of polymorphism in chromosomal regions affected by this force (Vekemans et al. 2021) it is still unclear how much of the genome is affected (Asthana et al. 2005; Fijarczyk and Babik 2015).

In addition to the sheer increase of polymorphism, balancing selection often promotes heterozygosity. As a consequence, deleterious mutations that are at least partially recessive may be sheltered from expression and purging in the linked regions. This has been proposed to diminish the efficacy of purifying selection and facilitate the local accumulation of a genetic load, referred to as the “sheltered load” (Uyenoyama 1997, 2005). The deterministic model by Leach et al. (1986) showed that very tight linkage is required for effective sheltering of lethal recessive mutations. More recently, Tezenas et al. (2023) observed that even though the deleterious recessive mutations partially linked to a fungal-like mating-type locus (where typically two strongly balanced allelic lines segregate) are almost surely purged, a limited fraction of them may persist for extended periods of time. Charlesworth et al. (1997) showed that sites linked to a locus under balancing selection evolve in a manner that is analogous to a geographically subdivided population, where each of the balanced allelic line has a small effective population size, and mildly deleterious linked mutations can rise to intermediate or high frequencies within them by genetic drift. Although this phenomenon would generate lineage-specific mutational load (Uyenoyama 1997), these deleterious mutations are generally kept rare by selection in the entire population, so the genetic load is expected to remain modest. Stochastic simulations by Llaurens et al. (2009) further predicted that the sheltering of fully linked deleterious recessive mutations should be related to the number of balanced allelic lines, with higher probabilities of fixation of deleterious alleles within an allelic line when the number of alleles increases. This possible accumulation of deleterious mutations has been considered as the “evolutionary cost” of balancing selection (Lenz et al. 2016), and in humans a large number of diseases are indeed associated with variants at genes linked to one of the classical examples of balancing selection in the human genome, the Major Histocompatibility Complex (*MHC*; e.g. Lenz et al. 2016; Matzaraki et al. 2017). In support of the sheltered load hypothesis, Lenz et al. (2016) observed a specific accumulation of putatively deleterious mutations (missense variants) in genes that are located inside the human *MHC* region but have no function in immunity and just happen to be linked to the *MHC* alleles. However, the important number of *HLA* genes under balancing selection were supposed to drastically restrict recombination in the *MHC* region. Beside this specific example, however, the effect of balancing selection on polymorphism of the linked chromosomal regions, in particular regarding the accumulation of a sheltered load, remains poorly documented.

Self-incompatibility (SI) in plants is perhaps the best understood case of long-term balancing selection (Castric and Vekemans 2004). This widespread genetic mechanism allows recognition and rejection of self-pollen, thereby preventing inbreeding and promoting outcrossing in hermaphrodites (Nettancourt 2001). Pollination between partners expressing identical haplotypes at the *S*-locus leads to rejection of the pollen, thus enforcing outcrossing and promoting higher heterozygosity than expected under random mating. In addition, as noted by Wright (1939), pollen produced by individuals carrying rare *S*-alleles will more rarely land on incompatible pistils than pollen produced by individuals carrying *S*-alleles that are more frequent. This results in strong negative frequency-dependent selection, which allows the stable maintenance of a large number of *S*-alleles within populations. Moreover, the *S*-alleles are maintained over very long evolutionary times (Vekemans and Slatkin 1994), and theoretical models predict that a local increase of nucleotide polymorphism should be observed in the linked genomic region (Uyenoyama 1997; Schierup et al. 2000). In gametophytic SI (GSI), pollen SI specificity is determined by its own haploid genome (as found e.g. in Solanaceae), whereas in sporophytic SI (SSI), the pollen recognition phenotype is determined by the male diploid parent (as found e.g. in Brassicaceae). In the Brassicaceae, SSI is controlled by a single genomic region, the *S*-locus (Schopfer, Nasrallah and Nasrallah 1999; Kusaba et al. 2001), composed of two linked genes, *SCR* (encoding the *S*-locus cysteine-rich protein) and *SRK* (encoding the *S*-locus receptor kinase protein), encoding the male and female specificity determinants, respectively.

The prediction that the *S*-locus could shelter a substantial genetic load that could influence the evolution of self-incompatibility systems (Uyenoyama 1997; Glémin et al. 2001) prompted a series of experimental studies to test the sheltered load hypothesis and quantify its magnitude, relying on controlled crosses to isolate the specific fitness effect of homozygosity at the *S*-locus. Such experiments have been performed in a number of plant species: *Papaver rhoea*s (Lane and Lawrence 1995), *Solanum carolinense* (Stone 2004), *Arabidopsis halleri* (Llaurens et al. 2009), *A. lyrata* (Stift et al. 2013) and *Rosa* (Vieira et al. 2021), revealing a measurable genetic load linked to the *S*-locus in several cases. The fact that this load was detectable at the phenotypic level, in spite of the inherently limited experimental power of these studies, suggests that the *S*-locus does indeed shelter a substantial load of deleterious mutations. However, a limitation is that these phenotypic studies could not evaluate the length of the chromosomal tract causing the observed phenotypic differences (potentially extending beyond the strictly linked regions), and thus provided no indication about the genomic architecture of the load. In *A. halleri* and *A. lyrata*, the *S*-locus has been sequenced entirely in multiple haplotypes, revealing that the non-recombining *S*-locus region contains no protein-coding genes besides the ones controlling the SI machinery itself (*SCR* and *SRK*; Guo et al. 2011; Goubet et al. 2012). The load detected phenotypically must therefore have been caused by mutations in the partially linked flanking regions rather than in the non-recombining *S*-locus region itself, which is puzzling given the prediction that only very tightly linked regions are expected to accumulate a substantial load. A series of empirical studies have set out to determine the genomic extent of the flanking region over which polymorphism is altered by linkage to the *S*-locus in *A. halleri* and *A. lyrata* (Kamau and Charlesworth 2005; Kamau et al. 2007; Ruggiero et al. 2008; Roux et al. 2013). These studies sequenced short fragments of a small subset of the genes immediately flanking the *S*-locus, as well as more distant genes, and compared their polymorphism to that of a handful of “control” coding sequences from across the genome. In these studies, the genomic extent of this increase was limited to the two first *S*-locus linked genes. However, the sparse sampling of genes and the sequencing of small gene fragments only did not allow these previous studies to reach solid conclusions on the true genomic extent of this increase, and to precisely evaluate the accumulation of “sheltered” deleterious mutations as compared to the genomic background.

In this study, we combined whole genome sequencing data with a targeted resequencing approach to comprehensively analyse all genes and intergenic sequences within 75kb on either side of the *S*-locus in three sample sets each of *A. halleri* and *A. lyrata*. We compared the observed patterns of polymorphism in these regions with those of 100 unlinked randomly chosen regions used as genomic controls. The use of internal genomic controls provides a powerful way to factor out differences in demographic histories and/or sample structure. We consistently observed an increase of total nucleotide polymorphism within the first 25-30 kb region immediately flanking the *S*-locus only, with no detectable effect further along the chromosome. However, this elevated overall polymorphism was not associated with a detectable increase of the frequencies at which putatively deleterious mutations segregate in these natural populations. Hence, even though linkage to the *S*-locus causes the accumulation of more total variation, including putatively deleterious mutations, recombination seems to be strong enough to prevent purifying selection from being effectively reduced even for the most tightly linked genes, or the high allelic diversity at the *S*-locus may prevent allele-specific linked mutations to reach high frequencies.

## Results

### Sequencing the S-locus flanking regions and control regions in large sample sets

To comprehensively evaluate the impact of balancing selection on the genomic regions flanking the *S-*locus, we re-sequenced the entirety of the chromosomal interval containing the *S*-locus over 75 kb on each side. We divided this region into three consecutive non-overlapping windows of 25kb (−25, −50 and −75kb on one side and +25, +50 and +75kb on the other side; Fig. 1). Together, these regions contained 33 annotated protein-coding genes in the *A. lyrata* genome (Hu et al. 2011), eleven of which were within the two 25kb windows closest to the *S*-locus, nine within the next upstream and downstream 25-50kb windows together, and thirteen within the most distant 50-75kb regions (Fig. 1). To compare these regions to the background level of nucleotide polymorphism, we also included in the analysis one hundred 25kb “control” regions unlinked to the *S*-locus. These control regions were randomly chosen across the *A. halleri* genome (Legrand et al. 2019) and selected to closely match the density of protein-coding sequences and transposable elements found at the *S*-locus flanking regions (proportion of CDS within the interval = 0.23 +/-0.0023, Fig. 1; proportion of TEs = 0.28 +/-0.0028). We also verified that the genes in the control regions had comparable lengths, number of introns and GC contents as the *S*-linked genes, Fig. S1). Because the extreme level of sequence divergence of the non-recombining interval containing the *S*-locus itself precludes mapping of short reads among *S*-haplotypes (Goubet et al. 2012), we excluded this region from further analysis and focused on the flanking regions only (Fig. 1).

**Figure 1:**
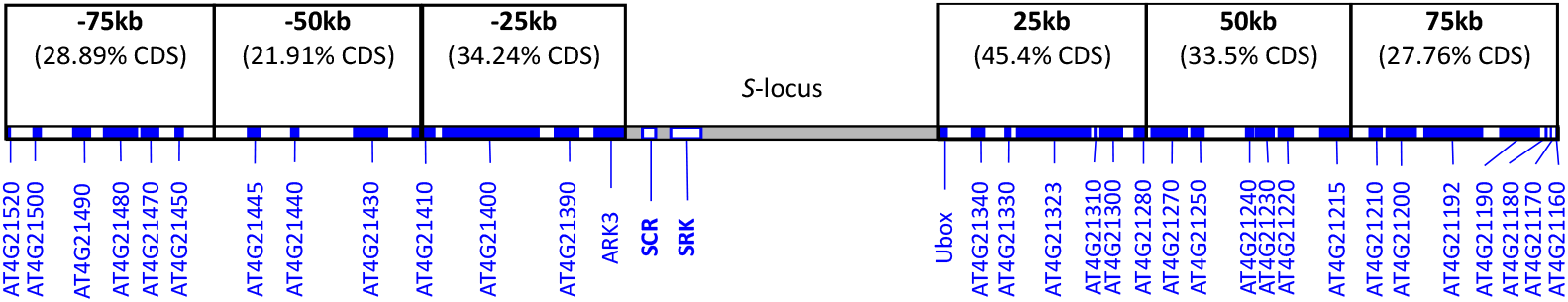
Schematic representation of the S-locus and its flanking regions. Protein-coding genes in the flanking regions are represented as filled blue rectangles. The genomic regions studied are distributed between positions 9,264,458 and 9,451,731 along chromosome 7 of the A. lyrata genome assembly (Hu et al. 2011). The S-locus (in grey) contains two protein-coding genes only, SRK and SCR (white rectangles) and is flanked by the ARK3 and Ubox genes (in the 5’ and 3’ directions, respectively). The S-locus region itself was not analysed in the present study. The percentage of CDS in each 25kb window is given on top of the figure.

To provide a comprehensive picture of the indirect effects of balancing selection, we analysed two closely related species that share the same orthologous SI system and show extensive trans-specific polymorphism at the *S*-locus, *A. halleri* and *A. lyrata* (Castric et al. 2008). In order to evaluate the robustness of our conclusions to different demographic histories, we analysed nucleotide sequence polymorphism from two natural populations of *A. halleri* (Nivelle, n=25, and Mortagne, n=27, that have been recently introduced in the North of France in association with industrial activities) and two natural populations of *A. lyrata* (Plech, n=18, from the core of the species range and Spiterstulen, n=23, from the edge of the species range, Table S1). To evaluate the robustness of our conclusions to different sampling strategies, we also included samples from more extended geographic regions of *A. lyrata* (North America, n=27, distributed across three distinct populations) and *A. halleri* ssp. *gemmifera* (Japan, n=47, distributed across six distinct populations). For the Nivelle, Mortagne and North American samples, we developed a dedicated sequence capture protocol specifically targeting the control and *S*-locus flanking regions. For the Japan, Plech and Spiterstulen samples, we took advantage of published whole-genome resequencing datasets, but analysed only polymorphism of the *S*-locus and control regions included in the capture protocol.

We obtained an average of 59 million reads mapped for the samples sequenced by sequence capture and 1,310 million reads for the WGS samples. After stringent filtering, we were able to interrogate with confidence an average of 960,368 positions in control regions, 48,014 of which were variable and biallelic (5%, Table S2). As expected, the number of variable sites across the control regions differed among sample sets, reflecting their different demographic histories (Table 1). The Japanese *A. halleri* sample set was the least polymorphic, with nucleotide polymorphism *π*=0.00128. At the other extreme, the *A. lyrata* Plech population was the most polymorphic, with *π*=0.00758. These estimations of the background level of nucleotide polymorphism in each sample set were used as internal genomic controls for the study of polymorphism in the *S-*flanking regions, where we were able to interrogate an average of 74,866 sites, containing 4,497 variable biallelic positions (6%, Table S2).

**Table 1:**
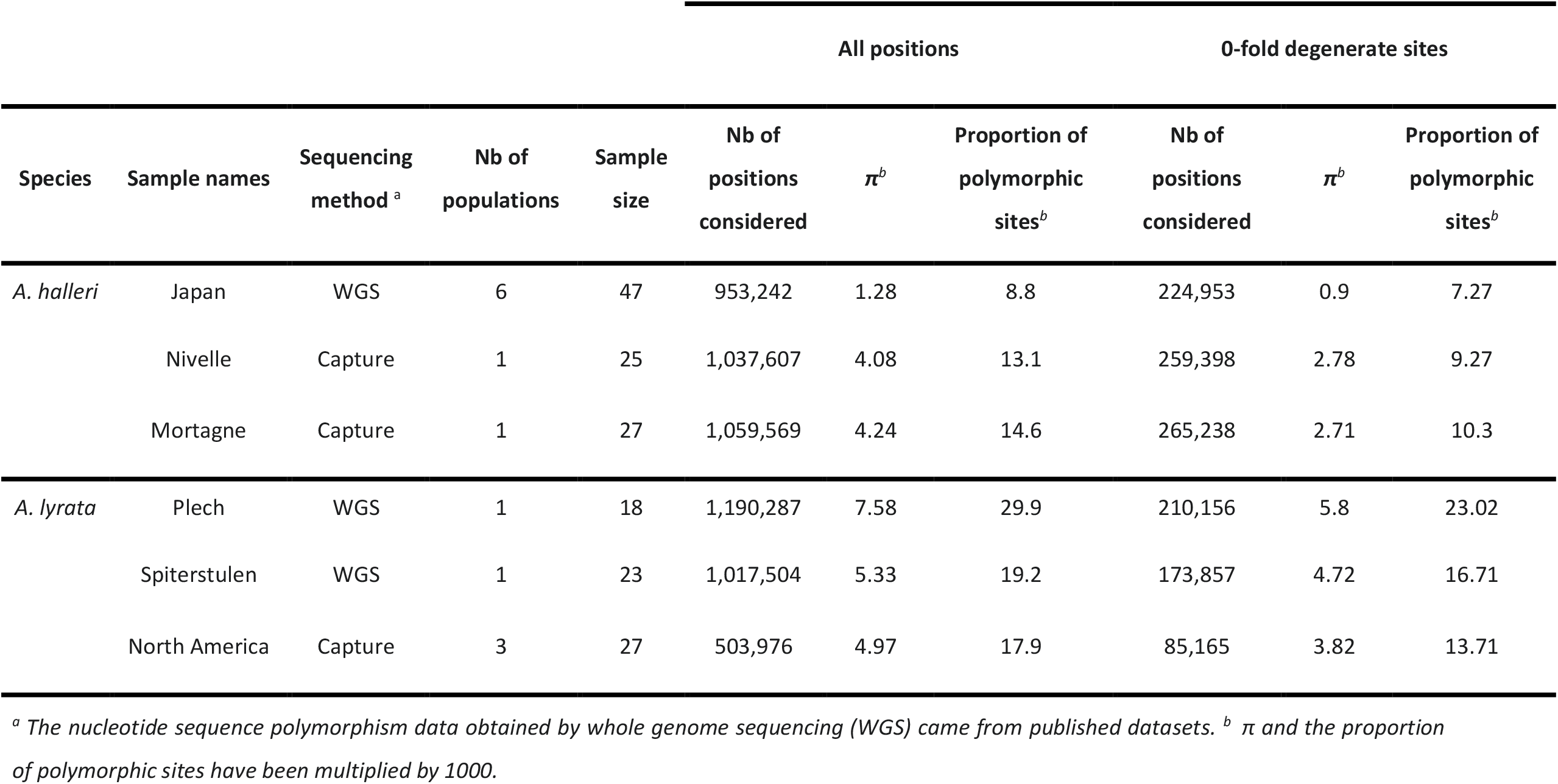
Variation of the median π and proportion of polymorphic sites in all positions and 0-fold degenerate sites in control regions in each dataset.

### Detection of the footprints of balancing selection on the S-linked regions

Based on these comprehensive polymorphism data, we combined different approaches to characterise the impact of balancing selection. As a first step, we excluded two potential confounding effects. First, we confirmed that the mean levels of raw divergence between the *A. thaliana* and *A. lyrata* reference genomes are similar between genes in the windows flanking the *S*-locus and those in the control regions, thus excluding the possibility that the linked genes simply accumulate more mutations per unit time. Instead, the two linked regions tended to exhibit a slight (non-significant) reduction rather than an increase of divergence (Fig. S2), making our analysis conservative. This observation is in line with the repeated introgression observed at the *S*-locus between *A. halleri* and *A. lyrata* (Castric et al. 2008), causing divergence between the two species to be more recent for the *S*-alleles than for the genomic background. Second, we used the recombination map of Hämäla et al. (2017), and confirmed that the recombination rate in the genomic interval containing the *S*-locus did not differ from those of the 100 control regions (Fig. S3).

These two confounding factors being excluded, we first followed a multilocus Hudson-Kreitman-Agade (HKA) approach to compare nucleotide polymorphism within *A. lyrata* and *A. halleri* sample sets, taking into account divergence from the outgroup *A. thaliana*. The test showed a significantly higher polymorphism of the *S*-flanking genes compared with the control loci (mean *k*=1.46, X^2^=821, df=33, *P*=0; Fig. S4, Tables S3-S5). This increase was more pronounced for linked genes closer to the *S*-locus, although the magnitude of this pattern varied across samples and also between the 5’ and 3’ flanking regions (Fig. S4, Tables S4 and S5).

We then used the new powerful approach of Cheng and Degiorgio (2020) that is robust to demographic variations to detect distortions of the site frequency spectrum along the chromosomal fragments and determine the maximum likelihood position of putative targets of balancing selection. We found signals of balancing selection in some of the control regions, particularly on chromosomes 3 and 4, but even though some of these signals were strong, they were not consistent across sample sets (Fig. 2). In contrast, the *S*-locus flanking regions carried consistent signals of balancing selection that were detected across all sample sets, even though the individual peak corresponding to the *S*-locus was not always the most extreme (Fig. 2). Overall, these results show that the *S*-locus stands out from the genomic background as the chromosomal location with the most consistent footprint of balancing selection.

**Figure 2:**
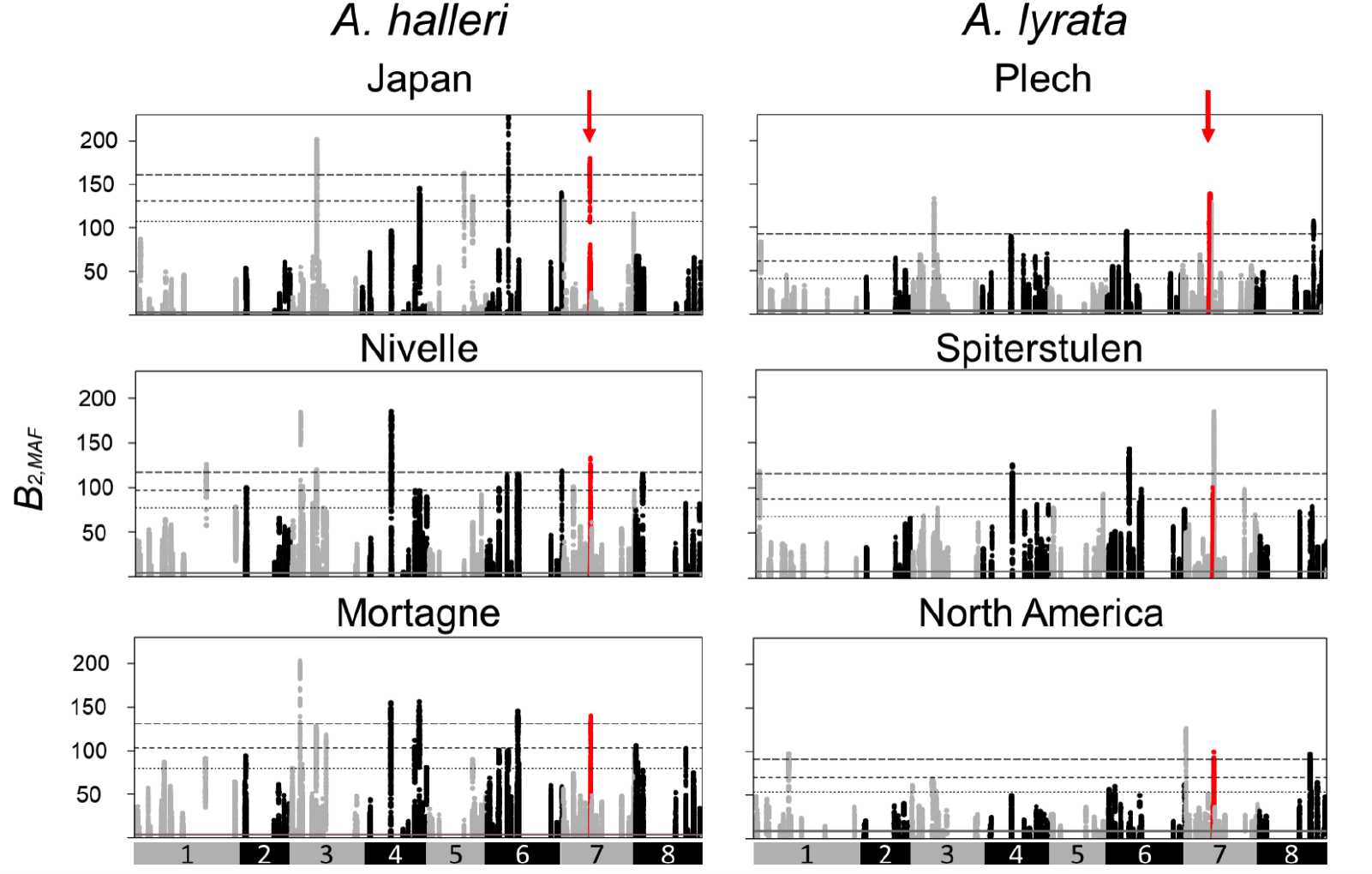
Manhattan plots for signals of balancing selection (B_2,MAF_ scores) in the 100 control regions (black and grey dots) and the S-locus region (red dots and arrows) along the A. lyrata genome. Chromosomes of the A. lyrata genome, with the 100 control regions distributed on successive chromosomes are represented by an alternation of grey and black dots. The horizontal grey solid line represents the median B_2,MAF_ scores across SNPs in the genome, and the black horizontal dashed lines represent the top 5, 2.5 and 1% percentiles.

### The increased polymorphism of the S-linked regions is mostly caused by an increase of the proportion of polymorphic sites

Then we sought to describe in more detail how polymorphism of the *S*-locus flanking regions differs from the genomic background. To do so, we compared the values of several summary statistics of polymorphism from the *S*-locus flanking regions to their distribution across the 100 control regions. A significant excess of total nucleotide polymorphism as compared to control regions was found for all sample sets in the two 25kb windows immediately flanking the *S*-locus (*π* was increased by a factor ranging from 1.6-fold in Plech to 5.8-fold in Japan; Fig. 3, Table S6). In stark contrast, the second and third consecutive 25kb windows on either side of the *S*-locus showed no excess polymorphism as compared to control regions in any sample set. We note that such a narrow genomic extent of the peak of nucleotide polymorphism is qualitatively consistent with predictions based on Takahata and Satta (1998)’s equations (Fig. S5) taking into account the local recombination rate deduced from the recombination map of Hämäla et al. (2017). To verify that the effect we observed was not specific to the particular window size we chose, we first repeated the comparison for the window immediately flanking the *S*-locus, increasing its size from 15kb to 75kb. The increased polymorphism was consistently observed when the window was below 30kb, but never when it was above 50kb (Fig. S6). Second, we repeated the comparison again, but on a gene-by-gene basis. Again, many of the genes with elevated nucleotide polymorphism were within the first 25kb window (Fig. S7A). Finally, to avoid defining arbitrary bins altogether, we used linear models to test whether *π* of individual sites of the *S*-flanking regions (considered as the response variable) declined with physical distance away from the *S*-locus. A highly significant negative effect of distance to the *S*-locus was observed overall (Table S7), confirming the effect of proximity to the *S*-locus on polymorphism of sites in the flanking regions.

**Figure 3:**
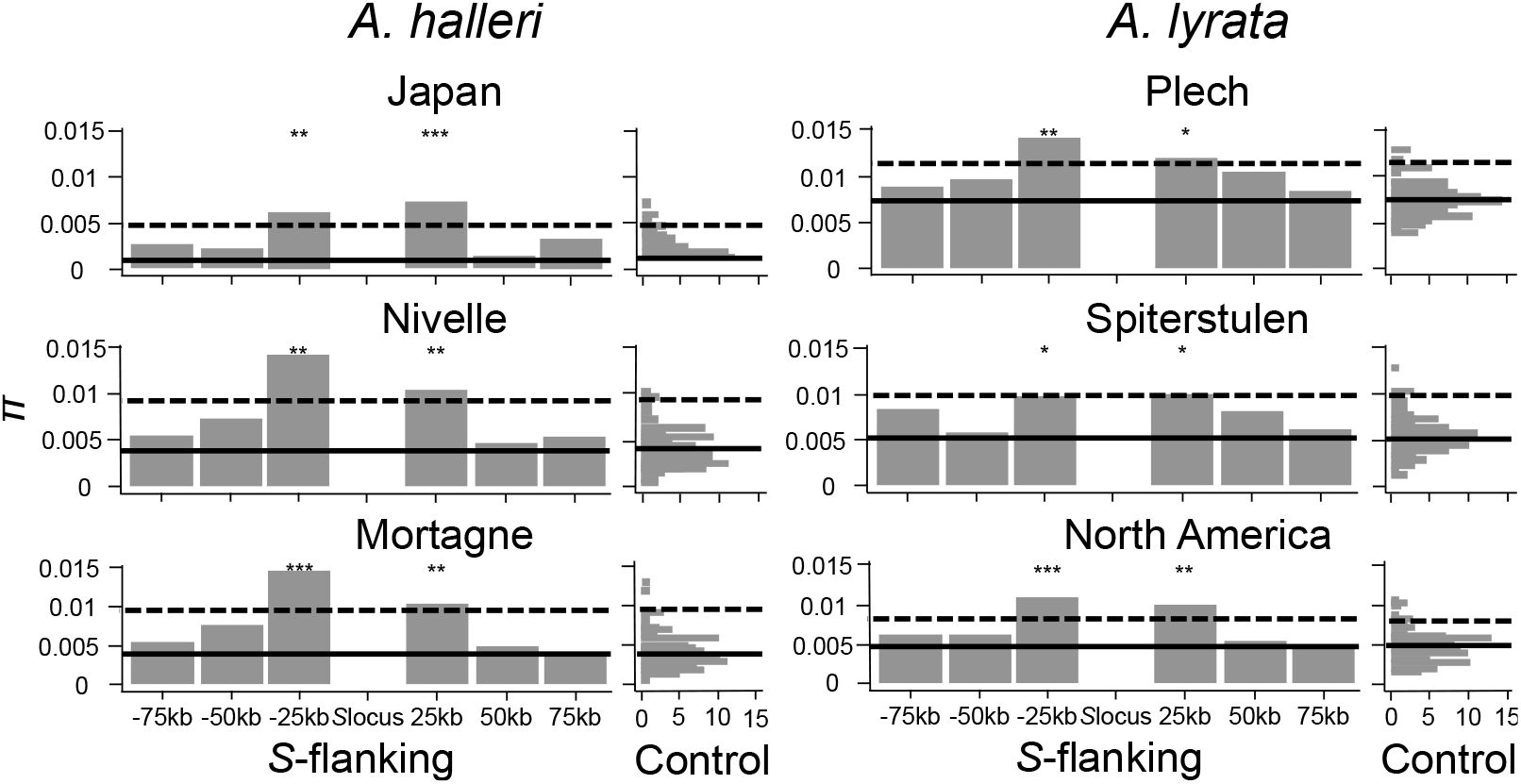
Mean π around the S-locus and across the control regions from throughout the genome. Each barplot represents the mean value of π obtained in non-overlapping regions of 25kb around the S-locus. The distributions (count) of π mean in the 100 control regions are represented by a vertical histogram on the right. The 95% percentile of the distributions is represented by dashed lines. The median value of the distribution in control regions is represented by black lines. *** = observed value above the 99% percentile of control regions, ** = observed value above the 97,5% of control regions, * = observed value above the 95% of control regions.

Then, we decomposed the nucleotide polymorphism (*π*) into its two components; *i*.*e*., the proportion of polymorphic sites (number of observed polymorphic sites divided by the total number of sites considered) and the mean frequency of the minor allele (*MAF*) at polymorphic sites. While the proportion of polymorphic sites was clearly increased relative to controls (by 1.6-fold in Plech to 3.9-fold in Japan, (Fig. S8A, Table S6), the *MAF* of polymorphic sites was no different than the genomic background (Fig. S8B, Table S6). Very similar patterns were observed when considering the 4-fold degenerate sites only (Table S8). Overall, as predicted, our results thus show elevated nucleotide polymorphism at the most-closely linked *S*-locus region, which is caused by a larger number of polymorphic sites rather than by an increased frequency at which the polymorphic sites segregate.

### No evidence for a weak purifying selection efficiency in the S-linked regions

To test whether the indirect effect of balancing selection described above is associated with the accumulation of a “sheltered” genetic load, we compared the polymorphism of 0-fold degenerate sites, assuming that the majority of amino-acid polymorphisms are deleterious to some extent (Eyre-Walker and Keightley 2007) to that of 4-fold degenerate sites. As for total and 4-fold degenerate polymorphisms above, we observed an increase of nucleotide polymorphism *π* at 0-fold degenerate sites, which was also predominantly due to an increased proportion of polymorphic sites in the first 25kb surrounding the *S*-locus (Table S9) but no change in *MAF*. The magnitude of these increases was comparable to that observed for total and 4-fold polymorphisms (Table S6 and S8 respectively), and a linear model restricted to 0-fold sites confirmed the effect of proximity to the *S*-locus (Table S7). In line with this observation and with predictions, the ratio *π*_0-fold_/*π*_4-fold_ in the *S*-locus flanking regions did not depart from its genomic distribution obtained from the controls, with the exception of the − 50kb window in the North American *A. lyrata* (Fig. 4). Hence, in spite of their predicted more pronounced effects on fitness, polymorphisms at the 0-fold degenerate sites did not tend to accumulate differently from those at 4-fold sites, suggesting that linkage to the *S*-locus was not associated with a decrease of purifying selection efficiency.

**Figure 4:**
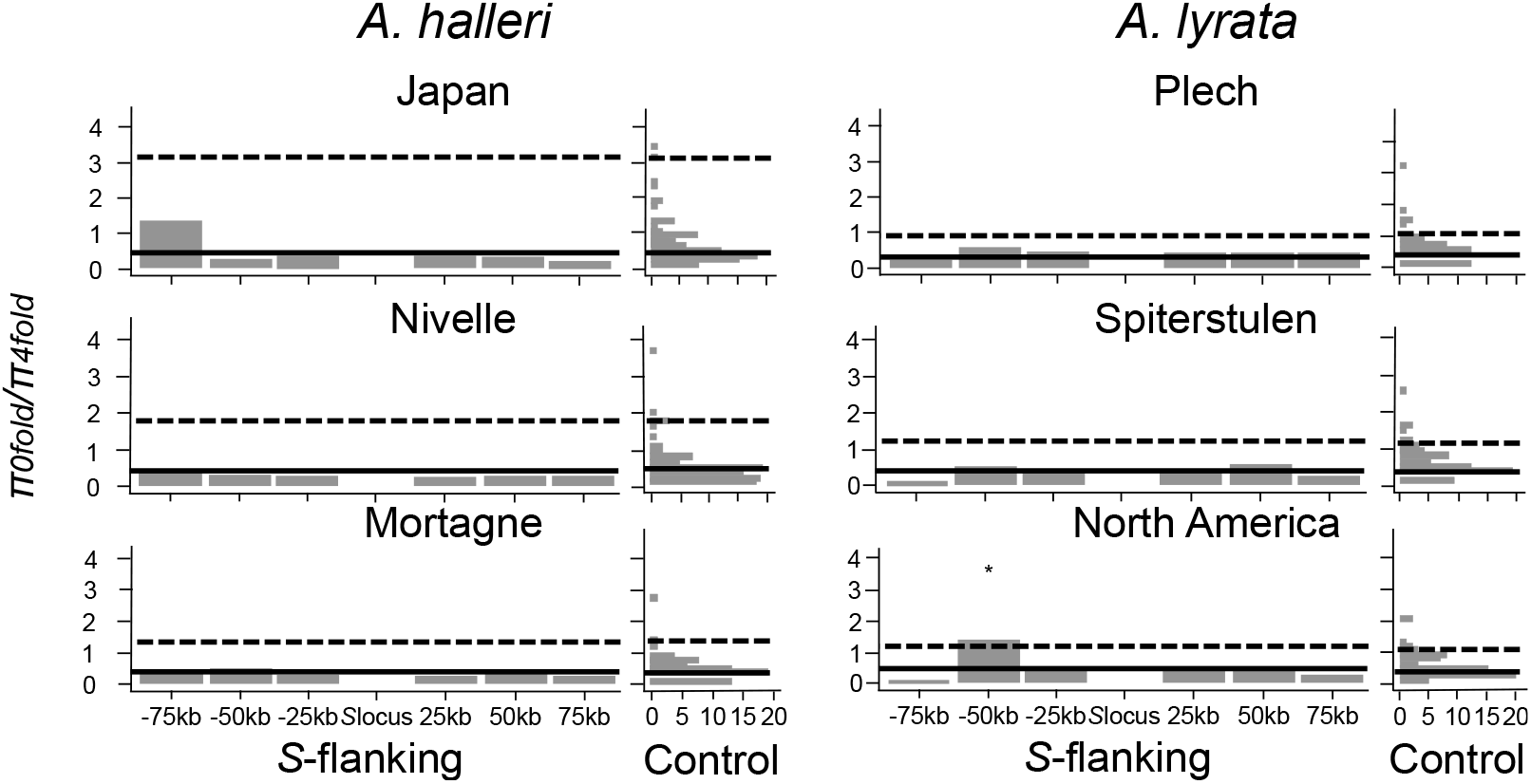
π_0-fold_/π_4-fold_ ratio around the S-locus and across the control regions from throughout the genome. The bars represent the π_0-fold_/π_4-fold_ ratio of sites in non-overlapping regions of 25kb around the S-locus. The distributions of the π_0-fold_/π_4-fold_ ratio in the 100 control regions are represented by a vertical histogram on the right. The 95% percentile of the distributions is represented by dashed lines. The median value of the distribution in control regions is represented by black lines. * = Observed value above the 95% percentile of control regions.

Finally, we explored the functional annotations of the eleven genes in the *S-*flanking regions where total nucleotide polymorphism was elevated (*i*.*e*. the two most tightly linked −25kb and +25kb intervals, Fig. 1). Four of these genes are receptor-like serine/threonine-protein kinases (*AT4G21410, AT4G21400, AT4G21390, AT4G21380*/*ARK3*), one is an ubiquitination protein (*AT4G21350*/*Ubox*), two are transcription factors (*AT4G21340, AT4G21330*), one a peptidase (*AT4G21323*), one a transmembrane protein (*AT4G21310*), one a tetratricopeptide repeat (TPR)-like superfamily protein (*AT4G21300*), and the last one is a subunit of Photosystem II (*AT4G21280*). Moreover, some individual genes further away from the *S*-locus had elevated polymorphism at 0-fold degenerate sites in several sample sets (Fig. S7B; Table S10), including a tetratricopeptide repeat (TPR)-like superfamily protein (*AT4G21170;* elevated for all populations with the exception of Japan and Spiterstulen), a DnaJ / Sec63 Brl domain-containing protein (*AT4G21180;* only elevated for the Japanese population), one cytochrome C oxidase (*AT4G21192;* elevated for the Mortagne and Nivelle populations), a LPxA enzyme involved in lipid metabolism (*AT4G21220;* elevated for the Mortagne, North America and Plech populations) and a F-box protein (*AT4G21240*; only elevated for the Plech population). Mutations at some of these genes were previously shown to be associated with important phenotypic traits such as the response to pathogens or biotic and abiotic stress, immunity, heterosis, root development, pollen sterility (Table S11).

## Discussion

### Confirmation of a limited extent of the genomic region affected by linkage to the S-locus

Our analysis across several replicate populations and geographic regions in two closely related plant species confirms that diversity is higher near the *S*-locus, as predicted, and reveals the extent of the genomic region affected. We find qualitatively very similar results across species and sampling regions, in spite of the different demographic histories, sampling structures and sequencing technologies used. However, we also note interesting quantitative differences. In particular, the relative elevation of polymorphism at the *S*-locus seems to be more pronounced in populations with lower baseline levels of diversity across the genome (the North American *A. lyrata* and the Japanese *A. halleri* samples). It would be interesting to determine whether the contrast is simply more readily detectable, or whether the strong genetic bottlenecks they experienced during their recent colonisation history (Clauss and Mitchell-Olds, 2006; Ross-Ibarra et al. 2008) may have decreased the number of *S*-alleles, leading to more intense balancing selection (Glémin et al. 2001, Billiard et al. 2007). Some North American populations have experienced a loss of SI, and have shifted to partial or complete selfing (Mable et al. 2005). Selfing is generally expected to reduce the effective rate of recombination, and might thus expand the footprint of balancing selection (Nordborg 1997). However, the populations considered here were specifically chosen because they are predominantly outcrossing (Foxe et al. 2010), and so we expect this effect to be minor. Interestingly, Hasselmann and Beye (2006) found similar level of genomic extent (<20 kb) of the region showing elevated polymorphism around the *csd* gene in honeybee, a gene subject to multiallelic balancing selection of the same order of magnitude as that acting on the S-locus (Hasselmann et al. 2008), although local recombination rates are higher in honeybee.

Our estimation of the size of the genomic region affected by linkage to the *S*-locus is largely in line with the theoretical prediction based on Takahata and Satta (1998)’s equations, although we caution that this comparison relies on a number of assumptions on the local rate of recombination, the number of *S*-alleles and their dominance interactions that each come with considerable uncertainty. First, while the *S*-locus seems to lie in a chromosomal interval whose recombination rate is no different from the genomic background, the recombination map available is still relatively coarse. Obtaining precise estimates of the local recombination rates around the *S*-locus thus will require denser genetic maps. Second, precisely estimating the number of *S*-alleles in a population remains challenging. While more reliable and higher throughput *S*-allele genotyping procedures have become available recently (Genete et al. 2020), the typically very large number of *S*-alleles segregating requires correspondingly large sample sizes. In our study, we decided to increase the number of populations compared and make comparisons across species, rather than focusing on larger sample sizes within a single population. Third, the species we studied (*A. halleri* and *A. lyrata*) have sporophytic SI, whereas the theoretical models predicting the effects of linkage are based on gametophytic SI or generic overdominance. Sporophytic systems are characterised by a complex dominance hierarchy (Prigoda et al. 2005; Llaurens et al. 2008), which would alter expectations for the strength of selection. For example, balancing selection acts more strongly on dominant *S*-alleles than on recessive *S*-alleles (Billiard et al. 2007), an important asymmetry that is not taken into account in current models. Furthermore, the dominance interactions between *S*-alleles have only been documented for a small set of the total number of possible pairs (Prigoda et al. 2005; Durand et al. 2014), and thus remain considerably uncertain.

### No evidence that linked genes experience a decrease of purifying selection efficiency

Our results suggest that the *S*-flanking regions do accumulate more potential deleterious mutations than the control regions, but these deleterious mutations segregate at population frequencies that are indistinguishable from those at control genes, and the *π*_*0-fold*_*/π*_*4-fold*_ ratio is also unchanged. Hence, the main factor causing the elevated polymorphism seems to be the deeper coalescence time among allelic lineages, allowing the accumulation of both neutral and deleterious mutations. Strikingly, our results are almost the mirror image of the pattern seen at the human *MHC*, where Lenz et al (2016) observed that the elevated polymorphism in the linked regions was caused by mutations that segregate at higher population frequencies than the genomic background, but not by an overall increased abundance of polymorphic sites across the region. For the *MHC*, balancing selection is believed to be driven by pathogen-mediated selection (although the exact mechanism remains controversial; see Spurgin and Richardson 2010), which is very different from the negative frequency-dependent selection maintaining diversity at the *S*-locus. Beside the fact that the *MHC* region is much larger and includes many more genes, it is currently not clear which specific feature of the balancing selection mechanism acting at the *S*-locus and the *MHC* causes these sharply distinct genomic signatures. The contrasted genomic effects detected in the *S*-flanking regions as compared to the MHC, reinforce the idea that the different evolutionary processes causing balancing selection can result in different genomic signatures.

One possible explanation for the fact that the distribution of allele frequencies in the linked regions are indistinguishable from those at the control regions can be related to the model proposed by Takahata (1990). This model shows that the genealogical relationships among distinct *S*-allele lineages under gametophytic SI are expected to be identical to those of neutral genes, except that they are expanded by a scaling factor *f*_S_. If the number of *S*-alleles in the studied sample sets was extremely large and every chromosome we sampled corresponded to a different *S*-allele lineage, then the only difference between the genealogies of sequences around the *S*-locus and those in the control regions would lie in the different time scales, hence overall polymorphism. Testing this hypothesis would require phasing the flanking sequences for each *S*-allele to obtain haplotypes. However, *S-*allele frequencies have been estimated in several of the population samples studied here (Llaurens et al. 2008; Genete et al. 2020; Takou et al. 2021), and it is already clear that randomly sampled chromosomes would be very unlikely to systematically correspond to distinct *S*-allele lineages. Nevertheless, theory predicts the possible fixation of a particular recessive deleterious allele only among gene copies associated with a given *S*-allele line, generating an allele-specific mutational load (Uyenoyama 2005), so that the population frequency of such deleterious alleles would remain low in populations with high *S*-allele diversity. In addition, the asymmetry introduced by the dominance interactions under the sporophytic system (driving recessive *S*-alleles to high population frequencies, whereas dominant *S*-alleles remain relatively rarer) could diminish the contrast between the *S*-locus and the genomic background as compared to the expectation under gametophytic SI.

### The identity of S-linked mutations

A series of previous studies aimed at experimentally revealing the phenotypic effect of *S*-linked mutations (Lane and Lawrence 1995; Stone 2004; Llaurens et al. 2009; Stift et al. 2013; Vieira et al. 2021). Such *S*-linked load affected different phenotypic traits across the different studies, as would be expected given that the *S*-locus lies in different genomic environments in distant species, and given that deleterious mutations are expected to hit different flanking genes in a random manner. In line with these observations, we also found that different genes exhibited an excess polymorphism at 0-fold degenerate sites as compared to the genomic background across the different populations analysed. Identifying more precisely the mutations causing these phenotypic effects would require fine mapping, and the present study indicates that they are most likely to be found in very close proximity to the *S*-locus (<25-30 kb). In fact, several of the genes found in this narrow chromosomal interval were previously shown to be associated with critical biological functions in plants such as photosynthesis (*AT4G21280*), abiotic and biotic stress responses (*AT4G21410; AT4G21400, AT4G21390, ARK3, AT4G21330* and *AT4G21300*), root development (*ARK3* and *AT4G21390*) or pollen fertility (*AT4G21330*). Notably, mutants of the transcription factor *AT4G21330* exhibit abnormal anther morphology at the beginning of stage four (Zhang et al. 2006) as well as magnesium-deficiency in leaves of *Citrus sinensis* (Ma et al. 2016) and pollen sterility in *Gossypium hirsutum* (Hamid et al. 2019). Hence, despite their limited number, some of these genes have important functions and are obvious candidates for the future dissection of the genetic architecture of the allele-specific mutation load sheltered by the *S*-locus.

In this context, a limitation of our population genetics approach is that it was designed to detect the collective accumulation of mutations rather than individual high-impact mutations. However, it is possible that a low number of high-impact mutations, rather than a collection of small-effect mutations, are causing the load. Indeed, the selective dynamics of lethal mutations *vs*. slightly deleterious mutations can be sharply different (e.g. Clo et al. 2020), and in the latter case finely dissecting the load at the genetic level will remain challenging. In addition, while our sequence capture approach also includes the intergenic sequences, we quantified the load based on coding sequences only. Previous studies demonstrated that polymorphism on intergenic regions could be under purifying selection (Mattila et al. 2019, for an example in *A. lyrata*), so it is also possible that besides the coding sequences, mutations in intergenic sequences contribute to the load, hence making our estimation of the sheltered load an underestimate. Structural variants in particular may have strong deleterious effects (Hämälä et al. 2021), but are entirely ignored in the present study. Long-read sequencing would be required to achieve a more detailed analysis of these types of polymorphisms.

An important difference between our population genetics approach and the previous phenotypic approaches is that here we integrate over all *S*-allele genotypic combinations present in our samples. In contrast, phenotypic studies are in practice limited to the analysis of a small number of *S*-alleles. For instance, Llaurens et al. (2009) and Stift et al. (2013) examined the effect of mutations linked to only three and four *S*-alleles respectively, inherently limiting the generality of their conclusions given the large number of *S*-alleles typically segregating in natural populations (Castric & Vekemans 2004). This strength of our approach is also a limitation, since we could not investigate the differential accumulation of mutations linked to different *S*-alleles. Llaurens et al. (2009) predicted a higher fixation probability of deleterious variants linked to dominant than to recessive *S*-alleles. Their phenotypic approach verified this prediction in *A. halleri*, but Stift et al. (2013) did not confirm it in *A. lyrata*. An exciting perspective to solve this discrepancy will be to compare the number and identity of deleterious mutations associated with dominant *vs*. recessive alleles, a task that will require phasing polymorphisms on a large number of *S*-linked haplotypes.

### Consequences for the (lack of) extension of the S-linked region

The segregation of deleterious mutations linked to the *S*-locus is expected to have important consequences for mating system evolution, affecting the maintenance of SI itself (Porcher and Lande 2005; Gervais et al. 2014) and the conditions for diversification of SI lineages (Uyenoyama 2003). As compared to other sex- or mating-type determining regions, however, the Brassicaceae *S*-locus has a remarkably limited size of the strictly linked genomic regions, containing no other gene beyond the SI determinants themselves. This is in strong contrast to the large chromosomal regions associated with sex-determining regions of many sex-chromosomes (Charlesworth 2017) or some mating-type loci in fungi (Hartmann et al. 2021). This cessation of recombination is believed to result from the progressive extension of successive inversions that can ultimately capture a large number of genes, eventually expanding across most of the length of a chromosome (Charlesworth et al. 2005; Otto et al. 2011). Recent models have shown that the successive fixation of inversions can take place even in the absence of sexual antagonism, as a result of the effective masking of recessive deleterious mutations accumulated in the flanking regions of the sex- or mating-type determining loci (Jay et al. 2022). We speculate that the limited size of the linked region upon which polymorphism is affected and the lack of effective sheltering by the *S*-locus, may contribute to prevent the region from expanding efficiently. It would now be interesting to investigate genomic patterns of the linked load in other SI systems, in particular in the non-self recognition SI system observed in Solanaceae, as the size of the *S*-locus is much larger in those systems (around 4Mb in a pericentromeric region of the *Petunia inflate* genome) and includes functional genes unrelated to SI (Wu et al. 2020).

## Material and methods

### Source plant material

We worked on natural accessions from two closely related species, *A. halleri* and *A. lyrata*, each represented by samples from three regions named Japan, Mortagne (France) and Nivelle (France) for *A. halleri*, and Plech (Germany), Spiterstulen (Norway) and North America for *A. lyrata* (Table S1). For the Japan, Spiterstulen and Plech samples, we used available whole genome sequencing (WGS) data obtained by Kubota et al. (2015) and Takou et al. (2021). The Japanese sample set was composed of 47 individuals of *A. halleri*, subsp. *gemmifera* originating from six different populations (17 individuals from Fujiwara, 17 from Ibuki, 2 from Inotani, 3 from Itamuro, 4 from Minoo and 4 from Okunikkawa; Kubota et al. 2015); the Spiterstulen (23 individuals) and Plech (18 individuals) sample sets were from single locations (Takou et al. 2021). For the three other sample sets, we collected individuals and developed a dedicated targeted enrichment capture approach to sequence the genomic regions of interest. The North American sample set of *A. lyrata* was composed of 26 individuals from three highly outcrossing populations from the Great Lakes region, named IND (Indiana Dunes National Lakeshore in Michigan, n=8), PIN (Pinery Provincial Park in Ontario, n=10) and TSS (Tobermory Provincial Park in Ontario, n=8) (Foxe et al. 2010). We collected 25 individuals from the Nivelle population (50°47’N, 3°47’E, France) and 27 individuals from the closely related Mortagne population (50°47’N, 3°47’E, France). In total, we complemented the 88 individuals with whole genome data available with an additional 78 newly sampled individuals that we sequenced with the targeted sequence capture approach described below.

### S-locus flanking regions and control regions

To evaluate the effect of balancing selection on the *S*-locus, we developed an original approach based on the comparison between the patterns of polymorphism of the two flanking regions on either side of the *S*-locus to those of a set of 100 randomly chosen control regions. The *S*-locus region can be poorly represented in whole-genome assemblies, so we first sequenced them using two *A. halleri* BAC clones that we newly obtained following the approach of Goubet et al. (2012) from a BAC library constructed from a mixture of several *A. halleri* individuals from Italy. These two BAC clones were chosen so as to cover entirely the 5’ and 3’ regions on either side of the *S*-locus (BAC clones 37G17 and 21E5, respectively; 10.6084/m9.figshare.16438908). We computed the proportion of coding sequences (CDS) and transposable elements (TEs) on the first 75kb sequences immediately flanking the *S*-locus on these two BAC clones (but excluding the non-recombining region within the *S*-locus itself), and we used these two statistics to select a set of matched control regions from across the *A. halleri* genome (Legrand et al. 2019). To do this, we used bedtools (Quinlan and Hall 2010) to randomly select 25 kb contiguous genomic intervals. The genomic intervals retained were characterised by a density of CDS and TEs closely matching that of the actual *S*-locus flanking regions (within 10%). The region with proportions of CDS and/or TEs departed from those values was discarded and a new region was picked until a total of 100 genomic intervals was included. The genomic coordinates of the control regions are given in Supplementary Table S12, and their sequences in fasta format are available at 10.6084/m9.figshare.16438908. Because the control regions were defined initially on the *A. halleri* reference, we used sequence similarity (using YASS, Noé and Kucherov 2005) to identify orthologous regions along the *A. lyrata* genome.

### Library preparation, sequence capture and sequencing

For the 78 newly sequenced individuals, we purified DNA from 15 mg of dried leaves of each sample with Chemagic beads (PerkinElmer) following Holtz et al. (2016), using the manufacturer’s instructions but with an additional Agencourt AMPure beads (Beckman) purification. DNA was quantified by Qubit and 50 ng of DNA was fragmented mechanically with Bioruptor (Diagenode) to obtain fragments of around 300bp, which we verified using a BioAnalyzer (Agilent) with a DNA HS chip. We prepared indexed genomic libraries using the Nextflex Rapid DNA Seq kit V2.0 (PerkinElmer) using the manufacturer’s instructions. Briefly, extremities of sequences were repaired and tailed, ligated with universal adaptors P5/P7 containing multiplexing unique dual index (PerkinElmer), and amplified by five cycles of PCR. We then selected fragments between 150 and 300pb with AMPures beads and pooled all libraries in equimolar proportions.

The pooled libraries then proceeded to a sequence capture protocol using the MyBaits v3 (Ann Arbor, Michigan, USA) approach. Briefly, 120bp RNA probes were designed by MyBaits and synthesised to target the complete set of one hundred 25kb control regions as well as the 75kb regions flanking the *S*-locus on either side, with an average tiling density of 2X (a total of 48,127 probes). In addition to the *S*-locus flanking regions and the control region, the capture array also contained a set of additional probes that were not used in the frame of the present project but are detailed in Supplementary Information. The indexed genomic libraries were hybridised to the probes overnight at a temperature of 65°C, and were finally sequenced by Illumina MiSeq (300bp, paired-end) by the LIGAN-MP Genomics platform (Lille, France).

### Read mapping and variant calling

Raw reads from sequence-capture or WGS datasets (see Supplementary table S1) were mapped onto the complete *A. lyrata* reference genome (V1.0.23, Hu et al. 2011) using Bowtie2 v2.4.1 (Langmead and Salzberg 2012). File formats were then converted to BAM using samtools v1.3.1 (Li et al. 2009) and duplicated reads were removed with the MarkDuplicates program of picard-tools v1.119 (http://broadinstitute.github.io/picard). These steps were performed by the custom Python script sequencing_genome_vcf.py available in https://github.com/leveveaudrey/analysis-of-polymorphism-*S*-locus. We retained only reads which mapped to the *S*-locus flanking or control regions. For the sake of consistency, we followed the same procedure for samples sequenced by WGS. Biallelic SNPs in these regions were called using the Genome Analysis Toolkit v. 3.8 (GATK, DePristo et al. 2011) with the option GVCF and a quality score threshold of 60 using vcftool v0.1.15 (Danecek et al. 2011). For each sample independently, we computed the distribution of coverage depth across control regions using samtools depth (Li et al. 2009). We excluded sites with either less than 15 reads aligned or coverage depth above the 97.5 % percentile, as the latter are likely to correspond to repeated sequences (e.g. transposable elements or paralogs). Sites covered by at least 15 reads but containing no SNP were considered as monomorphic. To exclude the possibility of spurious heterozygosity induced by the presence of paralogs, we verified that no SNP was systematically heterozygous across all individuals. The final number of sites in each sample set is summarised in Table 1. Finally, we excluded the sites covered by at least 15 reads but containing more than two SNPs (2.8%; table S2).

Because recombination rate is an important factor to explain variation of polymorphism, we used the recombination map of *A. lyrata* (Hämälä et al. 2017) to compare the recombination rate in *S*-flanking regions and in the control regions.

### Footprints of balancing selection

For each sample set, we first evaluated the distribution of the B_*2,MAF*_ statistic across all SNPs, which was designed to capture the distortion of the site frequency spectrum along chromosomes caused by linkage to a site under balancing selection (Cheng and Degiorgio 2020). We then compared the B_*2,MAF*_ distribution in control regions with the *S-*flanking regions, and considered a significant difference when the mean B_*2,MAF*_ value was outside the 95% percentile of the distribution in control regions.

To control for a possible difference in mutation rates between genes in the *S*-locus flanking regions and genes in the control regions, we then compared their pattern of molecular divergence between *A. lyrata* and *A. thaliana* (TAIR10 genome) at the sites retained for the polymorphism analysis (i.e. having passed the coverage filters). We identified orthologs as the best hits in the *A. thaliana* genome using YASS, retaining alignments with a maximum e-value of 0.01 and an identity above 70%. Pairs of sequences were then aligned with clustalOmega (Sievers et al. 2011) and the proportion of divergent sites was determined using a custom Python script (https://github.com/leveveaudrey/analysis-of-polymorphism-*S*-locus).

We further compared the ratio of within-species polymorphism to between-species divergence (Hudson et al. 1987) using the multilocus maximum likelihood HKA framework developed by Wright and Charlesworth (2004) and available at https://github.com/rossibarra/MLHKA. The algorithm is currently limited to only one hundred genes, so we tested the 33 *S*-locus flanking genes and a randomly chosen subset of *n*=67 of the 100 control genes. Specifically, we compared a model with free mutation at each locus and no selection against a model with free mutation but where each of the 33 *S*-locus flanking genes are allowed to have their own selection coefficient (*k*). This parameter corresponds to the relative increase of polymorphism of the *S*-linked genes compared to genes in the control regions, taking into account differences in divergence between *A. lyrata* and *A. thaliana* across loci. We used a log-likelihood ratio test with 33 degrees of freedom to compare the likelihood of these two nested models. Chain length was set to 100,000 and separate analyses were performed for each sample set independently. Before using the MLHKA framework, we compared the mean GC content (%) and the mean size of all exons cumulated by gene between the 33 *S*-flanking and 67 control genes by permutation tests (ten thousand reiterations). The mean size of all exons cumulated by gene was not significantly different between the *S*-flanking and control genes (p value=0.47). The mean size of all exons cumulated by gene was not significantly different between the *S*-flanking and control genes (p value=0.47). The mean GC content by gene was significantly higher in control genes compared with the *S*-flanking (+1.8, p value=0.01). This increase was explained by four *S*-flanking genes with a low GC content (Fig. S1), but no increase of polymorphism was detected on these genes.

### Decomposing the signals of balancing selection

We then decomposed the signal of balancing selection across the *S*-locus flanking regions into a series of elementary statistics. For each site, we estimated the minor allele frequency (*MAF*) and we calculated *π* at each position using the vcftools –site-pi option (Danecek et al. 2011). When a position of the *A. lyrata* genome was covered but not polymorphic, the *MAF* and *π* statistics were set to 0. For each statistic, we binned SNPs flanking the *S*-locus into 25kb intervals and compared the distribution of the mean value obtained for sites within non-overlapping windows of 25kb in the *S*-locus flanking regions with the distribution of the mean obtained across the 100 control regions. Moreover, we used General Linear Models (function *glmer* of the R package *lme4*, Bates et al. 2015) to test for a linear correlation between the exact distance of each SNP to the *S*-locus along the chromosome and each of the polymorphism statistics listed above, with the sample sets treated as a random effect.

We also compared the mean *π* obtained in the two *S*-flanking windows of different sizes (15, 25, 30, 35, 45, 55, 65 and 75kb) with the *π* obtained in the control regions in each dataset independently to define the minimum size of *S*-flanking regions with a significant excess of polymorphism compared with the genomic background. Then, we compared the mean *π* obtained in each gene with the *π* obtained in the control regions in each dataset independently to determine the genes in the *S*-flanking regions with a significant excess of polymorphism compared with the genomic background.

We also obtained a crude estimate of the linked region expected to show elevated neutral diversity around a balanced polymorphism by applying Takahata and Satta (1998)’s equations developed for a neutral locus partially linked to an overdominant locus, taking into account the local recombination rate in the *S*-locus region as deduced from the recombination map of Hämäla et al. (2017) (see Supplementary Information).

### Quantifying the sheltered load of deleterious mutations

To determine the extent to which the *S*-locus flanking regions accumulate deleterious mutations, we first reiterated the same analysis with the previous parameters (*MAF, π*), but for the 0-fold degenerate sites only (determined using the script NewAnnotateRef.py; Williamson et al. 2014). We assumed that all nonsynonymous changes are deleterious. Because all mutations at 0-fold degenerate sites alter the sequence of the encoded protein, we assumed that these mutations are deleterious (neglecting the rare cases where balancing selection could favour amino acid changes). In contrast, mutations at the 4-fold degenerate sites never alter the encoded amino acid, so we used them as neutral references. For the sake of simplicity, we discarded mutations at 2- or 3-fold degenerate sites.

## Acknowledgements

This work was supported by a grant from the France-Berkeley Fund (to VC and Rasmus Nielsen); the European Research Council (NOVEL project, grant number 648321); and the Agence Nationale de la Recherche (TE-MoMa project, grant number ANR-18-CE02-0020-01). AL thanks the ERC and the University of Lille for funding her PhD project. The authors thank the UMR 8199 LIGAN-MP Genomics platform (Lille, France), which belongs to the ‘Federation de Recherche’ 3508 Labex EGID (European Genomics Institute for Diabetes; ANR-10-LABX-46) and was supported by the ANR Equipex 2010 session (ANR-10-EQPX-07-01; ‘LIGAN-MP’). The LIGAN-PM Genomics platform (Lille, France) is also supported by the FEDER and the Region des Hauts-de-France. We thank Tuomas Hämälä for sharing the *A. lyrata* recombination map. We thank Stephen I. Wright, Rasmus Nielsen, Violaine Llaurens, Sylvain Glémin and Camille Roux for helpful discussions. The authors thank the Région Hauts-de-France, and the Ministère de l’Enseignement Supérieur et de la Recherche (CPER Climibio), and the European Fund for Regional Economic Development for their financial support for the molecular facilities in Lille. This work has been performed using infrastructure and technical support of the Plateforme Serre, cultures et terrains expérimentaux – Université de Lille for the greenhouse/field facilities.

Supplementary data include Fasta and Bed files of *A. halleri* regions and probes used for the sequence capture available online in figshare database at 10.6084/m9.figshare.16438908. All original sequence data are available in the NCBI Short Read Archive (SRA; https://www.ncbi.nlm.nih.gov/sra) with accession codes: PRJNA744343. All scripts developed are available in Github (https://github.com/leveveaudrey/analysis-of-polymorphism-S-locus).

## Bibliography

Aglyamova A, Petrova N, Gorshkov O, et al. 2022. Growing Maize Root: Lectins Involved in Consecutive Stages of Cell Development. Plants (Basel). 11:1799.

Asthana S, Schmidt S, Sunyaev S. 2005. A limited role for balancing selection. Trends in Genetics. 21: 30–32.

Bates D, Mächler M, Bolker B, Walker S. 2015. Fitting Linear Mixed-Effects Models Using lme4. Journal of Statistical Software. 67(1): 1–48.

Billiard S, Castric V, Vekemans, X. 2007. A general model to explore complex dominance patterns in plant sporophytic self-incompatibility systems. Genetics. 175: 1351–1369.

Castric V, Vekemans X. 2004. Plant self-incompatibility in natural populations: a critical assessment of recent theoretical and empirical advances. Molecular Ecology. 13: 2873–2889.

Castric V, Bechsgaard J, Schierup MH, Vekemans X. 2008. Repeated adaptive introgression at a gene under multiallelic balancing selection. PLoS Genetics. 4:e1000168.

Chae L, Sudat S, Dudoit S, et al. 2009. Diverse Transcriptional Programs Associated with Environmental Stress and Hormones in the Arabidopsis Receptor-Like Kinase Gene Family. Mol Plant. 2:84–107.

Charlesworth B, Morgan MT, Charlesworth D. 1993. The effect of deleterious mutations on neutral molecular variation. Genetics. 134: 1289–1303.

Charlesworth B, Nordborg M, Charlesworth D. 1997. The effects of local selection, balanced polymorphism and background selection on equilibrium patterns of genetic diversity in subdivided inbreeding and outcrossing populations. Genetics Research. 70: 155–174.

Charlesworth D, Charlesworth B, Marais G. 2005. Steps in the evolution of heteromorphic sex chromosomes. Heredity. 95: 118–128.

Charlesworth D. 2006. Balancing selection and its effects on sequences in nearby genome regions. PLoS Genetics. 2:e64.

Charlesworth D. 2017. Evolution of recombination rates between sex chromosomes. Philos Trans R Soc Lond B Biol Sci. 372(1736): 20160456.

Cheng X, DeGiorgio M. 2020. Flexible mixture model approaches that accommodate footprint size variability for robust detection of balancing selection. Molecular Biology and Evolution. 37: 3267–3291.

Clauss MJ, Mitchell-Olds T. 2006. Population genetic structure of Arabidopsis lyrata in Europe. Molecular Ecology. 15: 2753–2766.

Clo J, Ronfort J, Awad DA. 2020. Hidden genetic variance contributes to increase the short-term adaptive potential of selfing populations. Journal of Evolutionary Biology. 33: 1203–1215.

Danecek P, Auton A, Abecasis G, Albers CA, Banks E, DePristo MA, Handsaker RE, Lunter G, Marth GT, Sherry ST, et al. 2011. The variant call format and VCFtools. Bioinformatics. 27: 2156–2158.

DeGiorgio M, Lohmueller KE, Nielsen R. 2014. A model-based approach for identifying signatures of ancient balancing selection in genetic data. PLoS Genetics. 10, e1004561.

Delph LF, Kelly JK. 2014. On the importance of balancing selection in plants. New Phytologist. 201: 45–56.

DePristo MA, Banks E, Poplin RE, Garimella KV, Maguire JR, Hartl C, Philippakis AA, del Angel G, Rivas MA, Hanna M, et al. 2011. A framework for variation discovery and genotyping using next-generation DNA sequencing data. Nature Genetics. 43: 491–498.

Dwyer KG, Kandasamy MK, Mahosky DI, Acciai J, Kudish BI, Miller JE, Nasrallah ME, Nasrallah JB. 1994. A superfamily of S locus-related sequences in Arabidopsis: diverse structures and expression patterns. The Plant Cell. 6: 1829–1843.

Engelmann JC, Schwarz R, Blenk S, et al. 2008. Unsupervised Meta-Analysis on Diverse Gene Expression Datasets Allows Insight into Gene Function and Regulation. Bioinform Biol Insights. 2:265–280.

Eyre-Walker A, Keightley PD. 2007. The distribution of fitness effects of new mutations. Nature Reviews Genetics. 8: 610–618.

Fijarczyk A, Babik W. 2015. Detecting balancing selection in genomes: limits and prospects. Molecular Ecology. 24: 3529–3545.

Foxe JP, Stift M, Tedder A, Haudry A, Wright SI, Mable BK. 2010. Reconstructing origins of loss of self-incompatibility and selfing in North american Arabidopsis lyrata: a population genetic context. Evolution. 64: 3495–3510.

Genete M, Castric V, Vekemans X. 2020. Genotyping and De Novo Discovery of Allelic Variants at the Brassicaceae Self-Incompatibility Locus from Short-Read Sequencing Data. Molecular Biology and Evolution. 37: 1193–1201.

Gervais C, Awad DA, Roze D, Castric V, Billiard S. 2014. Genetic architecture of inbreeding depression and the maintenance of gametophytic self-incompatibility. Evolution 68, 3317–3324.

Goubet PM, Bergès H, Bellec A, Prat E, Helmstetter N, Mangenot S, Gallina S, Holl AC, Fobis-Loisy I, Vekemans X, et al. 2012. Contrasted patterns of molecular evolution in dominant and recessive self-incompatibility haplotypes in Arabidopsis. PLoS Genetics. 8, e1002495.

Glemin S, Bataillon T, Ronfort J, Mignot A, Olivieri I. 2001. Inbreeding Depression in Small Populations of Self-Incompatible Plants. Genetics. 159: 1217–1229.

Guo Y, Zhao X, Lanz C, Weigel D. 2011. Evolution of the S-locus region in Arabidopsis thaliana relatives. Plant Physiology. 157: 937–46.

Hämälä T, Mattila TM, Leinonen PH, Kuittinen H, Savolainen O. 2017. Role of seed germination in adaptation and reproductive isolation in Arabidopsis lyrata. Molecular Ecology. 26: 3484–3496.

Hämälä T, Wafula EK, Guiltinan MJ, Ralph PE, dePamphilis CW, Tiffin P. 2021. Genomic structural variants constrain and facilitate adaptation in natural populations of Theobroma cacao, the chocolate tree. PNAS 118 (35) e2102914118.

Hamid R, Marashi H, Tomar RS, et al. 2019. Transcriptome analysis identified aberrant gene expression in pollen developmental pathways leading to CGMS in cotton (Gossypium hirsutum L.). PLoS One. 14:e0218381.

Hartmann FE, Duhamel M, Carpentier F, Hood ME, Foulongne-Oriol M, Silar P, Malagnac F, Grognet P, Giraud T. 2021. Recombination suppression and evolutionary strata around mating-type loci in fungi: documenting patterns and understanding evolutionary and mechanistic causes. New Phytologist 229(5): 2470–2491.

Hasselmann M, Beye M. 2006. Pronounced differences of recombination activity at the sex determination locus of the honeybee, a locus under strong balancing selection. Genetics. 174(3):1469–80.

Hasselmann M, Vekemans X, Pflugfelder J, Koeniger N, Koeniger G, Tingek S, Beye M. 2008. Evidence for convergent nucleotide evolution and high allelic turnover rates at the complementary sex determiner gene of Western and Asian honeybees. Molecular Biology and Evolution. 25(4): 696–708.

Holtz Y, Ardisson M, Ranwez V, Besnard A, Leroy P, Poux G, Roumet P, Viader V, Santoni S, David J. 2016. Genotyping by sequencing using specific allelic capture to build a high-density genetic map of Durum Wheat. PLoS ONE. 11, e0154609.

Hu TT, Pattyn P, Bakker EG, Cao J, Cheng JF, Clark RM, Fahlgren N, Fawcett JA, Grimwood J, Gundlach H, et al. 2011. The Arabidopsis lyrata genome sequence and the basis of rapid genome size change. Nature Genetics. 43: 476–481.

Hudson RR, Kreitman M, Aguadé M. 1987. A test of neutral molecular evolution based on nucleotide data. Genetics. 116: 153–159.

Hudson RR, Kaplan NL. 1988. The coalescent process in models with selection and recombination. Genetics. 120: 831–840.

Jay P, Tezenas E, Véber A, Giraud T. 2022 Sheltering of deleterious mutations explains the stepwise extension of recombination suppression on sex chromosomes and other supergenes. PLoS Biology. 20(7):e3001698.

Kamau E, Charlesworth D. 2005. Balancing selection and low recombination affect diversity near the self-incompatibility loci of the plant Arabidopsis lyrata. Current Biology. 15: 1773–1778.

Kamau E, Charlesworth B, Charlesworth D. 2007. Linkage disequilibrium and recombination rate estimates in the self-incompatibility region of Arabidopsis lyrata. Genetics. 176: 2357–2369.

Kissen R, Overby A, Winge P, Bones AM. 2016. Allyl-isothiocyanate treatment induces a complex transcriptional reprogramming including heat stress, oxidative stress and plant defence responses in Arabidopsis thaliana. BMC Genomics. 17:740.

Kubota S, Iwasaki T, Hanada K, Nagano AJ, Fujiyama A, Toyoda A, Sugano S, Suzuki Y, Hikosak, K, Ito M, et al. 2015. A genome scan for genes underlying microgeographic-scale local adaptation in a wild arabidopsis species. PLoS Genetics. 11, e1005361.

Kusaba M, Dwyer K, Hendershot J, Vrebalov J, Nasrallah JB, Nasrallah ME. 2001. Self-incompatibility in the genus Arabidopsis: characterization of the S locus in the outcrossing A. lyrata and its autogamous relative A. thaliana. The Plant Cell. 13: 627–643.

Lane MD, Lawrence MJ. 1995. The population genetics of the self-incompatibility polymorphism in Papaver rhoeas. X. An association between incompatibility genotype and seed dormancy. Heredity. 75: 92–97.

Langmead B, Salzberg SL. 2012. Fast gapped-read alignment with Bowtie 2. Nature Methods. 9: 357–359.

Lauss K, Wardenaar R, Oka R, van Hulten MHA., Guryev V, Keurentjes JJB., Stam M, Johannes F. 2018. Parental DNA Methylation States Are Associated with Heterosis in Epigenetic Hybrids. Plant Physiol. 176: 1627–1645.

Lavagi-Craddock I, Dang T, Comstock S, et al. 2022. Transcriptome Analysis of Citrus Dwarfing Viroid Induced Dwarfing Phenotype of Sweet Orange on Trifoliate Orange Rootstock. Microorganisms 10:1144.

Leach CR, Mayo O, Morris MM. 1986. Linkage disequilibrium and gametophytic self-incompatibility. Theoretical and Applied Genetics. 73: 102–112.

Legrand S, Caron T, Maumus F, Schvartzman S, Quadrana L, Durand E, Gallina S, Pauwels M, Mazoyer C, Huyghe L, et al. 2019. Differential retention of transposable element-derived sequences in outcrossing Arabidopsis genomes. Mobile DNA. 10: 30.

Lenz TL, Spirin V, Jordan DM, Sunyaev S.R. 2016. Excess of deleterious mutations around HLA genes reveals evolutionary cost of balancing selection. Molecular Biology and Evolution. 33: 2555–2564.

Li H, Handsaker B, Wysoker A, Fennell T, Ruan J, Homer N, Marth G, Abecasis G, Durbin R, 1000 Genome Project Data Processing Subgroup. 2009. The Sequence Alignment/Map format and SAMtools. Bioinformatics. 25: 2078–2079.

Llaurens V, Billiard S, Leducq JB, Castric V, Klein E.K, Vekemans X. 2008. Does frequency-dependent selection with complex dominance interactions accurately predict allelic frequencies at the self-incompatibility locus in Arabidopsis halleri? Evolution. 62: 2545–2557.

Llaurens V, Gonthier L, Billiard S. 2009. The sheltered genetic load linked to the S locus in plants: new insights from theoretical and empirical approaches in sporophytic self-incompatibility. Genetics. 183: 1105–1118.

Ma CL, Qi YP, Liang WW, et al. 2016. MicroRNA Regulatory Mechanisms on Citrus sinensis leaves to Magnesium-Deficiency. Front Plant Sci. 7:201.

Mable BK, Robertson AV, Dart S, Di Berardo C, Witham L. 2005. Breakdown of self-incompatibility in the perennial Arabidopsis lyrata (Brassicaceae) and its genetic consequences. Evolution. 59(7): 1437–48.

Martins ACQ, Mota APZ, Carvalho Pasv, et al. 2022. Transcriptome Responses of Wild Arachis to UV-C Exposure Reveal Genes Involved in General Plant Defense and Priming. Plants (Basel). 11:408.

Mattila TM, Laenen B, Horvath R, Hämälä T, Savolainen O, Slotte T. 2019. Impact of demography on linked selection in two outcrossing Brassicaceae species. Ecology and Evolution. 9: 9532–9545.

Matzaraki V, Kumar V, Wijmenga C, Zhernakova A. 2017. The MHC locus and genetic susceptibility to autoimmune and infectious diseases. Genome Biology. 18: 76.

Meier S, Ruzvidzo O, Morse M, et al. 2010. The Arabidopsis Wall Associated Kinase-Like 10 Gene Encodes a Functional Guanylyl Cyclase and Is Co-Expressed with Pathogen Defense Related Genes. PLoS One. 5:e8904.

Mondal R, Biswas S, Srivastava A, Basu S, Trivedi M, Singh SK, Mishra Y. 2021. In silico analysis and expression profiling of S-domain receptor-like kinases (SD-RLKs) under different abiotic stresses in Arabidopsis thaliana. BMC Genomics. 22: 817.

Nettancourt D. 2001. Incompatibility and incongruity in wild and cultivated plants. Berlin Heidelberg: Springer-Verlag.

Noé L, Kucherov G. 2005. YASS: enhancing the sensitivity of DNA similarity search. Nucleic Acids Research. 33: W540–W543.

Noman M, Jameel A, Qiang WD, et al. 2019 Overexpression of GmCAMTA12 Enhanced Drought Tolerance in Arabidopsis and Soybean. Int J Mol Sci. 20:4849.

Nordborg M. 1997. Structured Coalescent Processes on Different Time Scales. Genetics 146: 1501–1514.

Otto SP, Pannell JR, Peichel CL, Ashman TL, Charlesworth D, Chippindale AK, Delph LF, Guerrero RF, Scarpino SV, McAllister BF. 2011. About PAR: The distinct evolutionary dynamics of the pseudoautosomal region. Trends in Genetics. 27: 358–367.

Pastuglia M, Swarup R, Rocher A, Saindrenan P, Roby D, Dumas C, Cock JM. 2002. Comparison of the expression patterns of two small gene families of S gene family receptor kinase genes during the defence response in Brassica oleracea and Arabidopsis thaliana. Gene. 282: 215–225.

Porcher E, Lande R. 2005. Loss of gametophytic self-incompatibility with evolution of inbreeding depression. Evolution 59, 46–60.

Prigoda NL, Nassuth A, Mable BK. 2005. Phenotypic and genotypic expression of self-incompatibility haplotypes in Arabidopsis lyrata suggests unique origin of alleles in different dominance classes. Molecular Biology and Evolution. 22(7): 1609–20.

Quinlan AR, Hall IM. 2010. BEDTools: a flexible suite of utilities for comparing genomic features. Bioinformatics. 26: 841–842.

Ross-Ibarra J, Wright SI, Foxe JP, Kawabe A, DeRose-Wilson L, Gos G, Charlesworth D, Gaut BS. 2008. Patterns of polymorphism and demographic history in natural populations of Arabidopsis lyrata. PLoS One 3, e2411.

Roux C, Pauwels M, Ruggiero MV, Charlesworth D, Castric V, Vekemans X. 2013. Recent and ancient signature of balancing selection around the S-locus in Arabidopsis halleri and A. lyrata. Molecular Biology and Evolution. 30: 435–447.

Ruggiero MV, Jacquemin B, Castric V, Vekemans X. 2008. Hitch-hiking to a locus under balancing selection: high sequence diversity and low population subdivision at the S-locus genomic region in Arabidopsis halleri. Genet Res (Camb). 90: 37–46.

Schierup MH, Vekemans X, Charlesworth D. 2000. The effect of hitch-hiking on genes linked to a balanced polymorphism in a subdivided population. Genetics Research. 76: 63–73.

Schierup MH, Mikkelsen AM, Hein J. 2001. Recombination, balancing selection and phylogenies in MHC and self-incompatibility genes. Genetics. 159: 1833–1844.

Schopfer CR, Nasrallah ME, Nasrallah JB. 1999. The male determinant of self-incompatibility in Brassica. Science. 286: 1697–1700.

Siever F, Wilm A, Dineen D, Gibson TJ, Karplus K, Li W, Lopez R, McWilliam H, Remmert M, Söding J, et al. 2011. Fast, scalable generation of high-quality protein multiple sequence alignments using Clustal Omega. Mol Syst Biol. 7: 539.

Smith JM, Haigh J. 1974. The hitchhiking effect of a favorable gene. Genetics Research. 23:23–35.

Spurgin LG, Richardson DS. 2010. How pathogens drive genetic diversity: MHC, mechanisms and misunderstandings. Proceedings of the Royal Society B: Biological Sciences. 277: 979–988.

Stift M, Hunter BD, Shaw B, Adam A, Hoebe PN, Mable BK. 2013. Inbreeding depression in self-incompatible North-American Arabidopsis lyrata: disentangling genomic and S-locus-specific genetic load. Heredity. 110: 19–28.

Stone JL. 2004. Sheltered load associated with S-alleles in Solanum carolinense. Heredity. 92: 335–342.

Strobeck C. 1980. Heterozygosity of a neutral locus linked to a self-Incompatibility locus or a balanced lethal. Evolution. 34: 779–788.

Strobeck C. 1983. Expected linkage disequilibrium for a neutral locus linked to a chromosomal arrangement. Genetics. 103: 545–555.

Takahata N. 1990. A simple genealogical structure of strongly balanced allelic lines and trans-species evolution of polymorphism. Proceedings of the National Academy of Sciences. 87: 2419–2423.

Takahata N, Nei M. 1990. Allelic genealogy under overdominant and frequency-dependent selection and polymorphism of major histocompatibility complex loci. Genetics. 124: 967–978.

Takahata N, Satta Y. 1998. Footprints of intragenic recombination at HLA loci. Immunogenetics. 47: 430–441.

Takou M, Hämälä T, Koch EM, Steige K.A, Dittberner H, Yant L, Genete M, Sunyaev S, Castric V, Vekemans X, et al. 2021. Maintenance of adaptive dynamics and no detectable load in a range-edge outcrossing plant population. Molecular Biology and Evolution. 38:1820–1836.

Tezenas E, Giraud T, Véber A, Billiard S. 2023. The fate of recessive deleterious or overdominant mutations near mating-type loci under partial selfing. Peer Community Journal. 3.

Uyenoyama MK. 1997. Genealogical structure among alleles regulating self-incompatibility in natural populations of flowering plants. Genetics. 147: 1389–1400.

Uyenoyama, MK. 2003. Genealogy-dependent variation in viability among self-incompatibility genotypes. Theoretical Population Biology. 63: 281–293.

Uyenoyama MK. 2005. Evolution under tight linkage to mating type. New Phytol. 165: 63–70.

Vekemans X, Slatkin M. 1994. Gene and allelic genealogies at a gametophytic self-incompatibility locus. Genetics. 137:1157–1165.

Vekemans X, Castric V, Hipperson H, Müller NA, Westerdahl H, Cronk Q. 2021. Whole-genome sequencing and genome regions of special interest: Lessons from major histocompatibility complex, sex determination, and plant self-incompatibility. Mol Ecol. 30: 6072–6086.

Vieira J, Pimenta J, Gomes A, Laia J, Rocha S, Heitzler P, Vieira CP. 2021. The identification of the Rosa S-locus and implications on the evolution of the Rosaceae gametophytic self-incompatibility systems. Sci Rep. 11: 3710.

Welgemoed T, Pierneef R, Sterck L, et al. 2020. De novo Assembly of Transcriptomes From a B73 Maize Line Introgressed With a QTL for Resistance to Gray Leaf Spot Disease Reveals a Candidate Allele of a Lectin Receptor-Like Kinase. Front Plant Sci. 11:191.

Williamson RJ, Josephs EB, Platts AE, Hazzouri KM, Haudry A, Blanchette M, Wright SI. 2014. Evidence for widespread positive and negative selection in coding and conserved noncoding regions of Capsella grandiflora. PLOS Genetics 10, e1004622.

Wiuf C, Zhao K, Innan H, Nordborg M. 2004. The probability and chromosomal extent of trans-specific polymorphism. Genetics. 168: 2363–2372.

Wright S. 1939. The distribution of self-sterility alleles in populations. Genetics 24: 538–552.

Wright SI, Charlesworth B. 2004. The HKA test revisited: a maximum likelihood ratio test of the standard neutral model. Genetics. 168: 1071–1076.

Wu L, Williams JS, Sun L, Kao TH. 2020. Sequence analysis of the Petunia inflata S-locus region containing 17 S-Locus F-Box genes and the S-RNase gene involved in self-incompatibility. The Plant Journal. 104: 1348–1368.

Yadeta KA, Elmore JM, Creer AY, et al. 2017. A Cysteine-Rich Protein Kinase Associates with a Membrane Immune Complex and the Cysteine Residues Are Required for Cell Death1[OPEN]. Plant Physiol. 173:771–787.

Zhang W, Sun Y, Timofejeva L, Chen C, Grossniklaus U, Ma H. 2006. Regulation of Arabidopsis tapetum development and function by DYSFUNCTIONAL TAPETUM1 (DYT1) encoding a putative bHLH transcription factor. Development. 133: 3085–3095.

Zhang W, Forester NT, Moon CD, et al. 2022. Epichloë seed transmission efficiency is influenced by plant defense response mechanisms. Frontiers in Plant Science. 13.

