## Supplementary data for "Long-term balancing selection and the genetic load linked to the self-incompatibility locus in *Arabidopsis halleri* and *A. lyrata*"

### Supplementary information

#### Capture array

The capture array was initially designed to enable the study of a range of genomic regions addressing different scientific questions in our lab. Thus, in addition to the *S*-locus flanking and control regions used in the present study, the capture array also contained a set of probes targeted towards: 1) a library of complete *S*-locus sequences for 36 *S*-alleles obtained from BAC clones ; 2) a library of 123 partial or complete *SRK* sequences from several brassicaceae species (*A. halleri, A. lyrata, A. thaliana, A. kamchatica, Capsella grandiflora, C. rubella, Brassica rapa, B. oleaceae*) ; 3) a set of 185 microRNA genes and their predicted mRNA target sites across the *A. haller*i genome; and 4) a candidate QTL region for heavy-metal tolerance. These additional probes were not used in the frame of the present project, and based on the absence of substantial sequence similarity, they are unlikely to interfere with our results. The complete list and sequences of the probes are available on the figshare database (10.6084/m9.figshare.16438908).

**Estimation of the size of the region flanking a balanced polymorphism that would show elevated neutral diversity.**

We obtained a crude estimate of the linked region expected to show elevated neutral diversity around a balanced polymorphism by applying Takahata and Satta (1998)'s equations developed for a neutral locus partially linked to an overdominant locus.
We adjusted their model's parameters in order to obtain a number of balanced allelic lines close to observed values in *A. halleri* (n = 19 in Japanese populations, Genete et al. 2021), and to obtain a background neutral polymorphism close to observed values ($\pi$ = 0.0025 in Japanese populations). We used the estimate of local recombination rate in the S-locus flanking region as deduced from the recombination map of Hämäla et al. (2017, r = 2.5 cM/Mb).

From equation (5) of Takahata & Satta (1998), the mean coalescence time of two genes randomly sampled from a population is given by:
$T = \frac{1}{n} T_w + \left(1 - \frac{1}{n}\right) T_b$, with *n* = number of balanced allelic lines, $T_w$ = mean coalescence time within an allelic line, $T_b$ = mean coalescence time between allelic lines.
*n* is given by their equation (2) : $n = \sqrt{\sqrt{2}S\left(\frac{u}{a}\right)}$, with $S = 2N_e s$ ($N_e$ is the effective population size, *s* is the coefficient of selection, set to its maximum value = 1), *u* is the rate of introduction of new allelic lines, and *a* is the rate of allelic turnover per line.
*a* is derived by their equation (1) : $a = \frac{u}{\sqrt{2}} ln\left(\frac{S}{16\pi M^2}\right)$, with $M = N_e u$.
$T_w$ is given by their equation (3): $T_w = \frac{2N_e(a+c^*n)}{(n+2N_e a)(a+c^*)}$, with $c^* = c\left(1 - \frac{1}{n}\right)$ where *c* is the recombination rate between the selected locus and the neutral locus.
$T_b$ is given by their equation (4): $T_b = T_w + \frac{n-1}{2(a+c^*)}$
Then, the nucleotide diversity at a given site partially linked to the selected locus, $\pi$, is given by: $\pi = 2T\mu$, with $\mu$ the mutation rate at the neutral locus.

As stated above, we adjusted the parameters to roughly mimic the balanced polymorphism at the S-locus. We set $N_e$ = 2.500, the rate of introduction of new selected alleles at $1.10^{-9}$, and the coefficient of selection *s* =1, which generates a number of balanced allelic lines close to *n* = 18. We then used $\mu = 1.5\ 10^{-7}$. which produces a nucleotide diversity of about 0.0025 in regions unlinked to the selected locus. Then we applied a local recombination rate in the flanking regions of the S-locus equal to 2.5 cM/Mb, as deduced from the recombination map of Hämäla et al. (2017).

Hämälä T, Mattila T.M, Leinonen P.H, Kuittinen H, Savolainen O. 2017. Role of seed germination in adaptation and reproductive isolation in *Arabidopsis lyrata*. Molecular Ecology. 26: 3484–3496.

Takahata N, Satta Y. 1998. Footprints of intragenic recombination at HLA loci. Immunogenetics. 47: 430–441.

**Supplementary data**

Tables

***Table S1: Summary of the sample sets used.***

| Species | Populations | Reference | Sample Size | Accession N° |
|---|---|---|---|---|
| *A. halleri* | Japan | *Kubota et al. 2015* | 47[a] | DRA003268 |
| | Nivelle | *This study* | 25 | PRJNA744343 |
| | Mortagne | *This study* | 27 | PRJNA744343 |
| *A. lyrata* | Plech | *Takou et al. 2021* | 18 | PRJEB34247, PRJEB33206 |
| | Spiterstulen | *Takou et al. 2021* | 23 | PRJEB34247, PRJEB33206 |
| | North America | *This study* | 26[b] | PRJNA744343 |

*[a] The sample of Japan was composed of 17 individuals from Fujiwara, 17 from Ibuki, 2 from Inotani, 3 from Itamuro, 4 from Minoo and 4 from Okunikkawa. [b] The sample of North America was composed of 8 individuals from IND, 10 from PIN and 8 from TSS.*

***Table S2: Number of nucleotide polymorphisms called across all control and S-flanking regions in each sample set.***

| | | **Control regions** | | ***S*-flanking regions** | |
|---|---|---|---|---|---|
| **Species** | **Population** | **Number of biallelic sites** | **More than two variants (excluded)** | **Number of biallelic sites** | **More than two variants (excluded)** |
| *A. halleri* | Japan | 17,092 | 395 | 2,060 | 74 |
| | Nivelle | 27,450 | 808 | 3,039 | 134 |
| | Mortagne | 28,914 | 932 | 3,151 | 131 |
| *A. lyrata* | Plech | 39,427 | 1,752 | 3,801 | 218 |
| | Spiterstulen | 22,738 | 508 | 2,775 | 91 |
| | North America | 9,559 | 211 | 1,604 | 54 |

***Table S3: Log likelihood-ratio obtained by the MLHKA test comparing the 33 S-flanking genes to a random set of 67 control genes .***

| Species | Population | Log likelihood-ratio | *P* value[a] |
|---|---|---|---|
| *A. halleri* | Japan | 1230 | 0 |
| | Nivelle | 750 | 0 |
| | Mortagne | 598 | 0 |
| *A. lyrata* | Plech | 422 | 0 |
| | Spiterstulen | 683 | 0 |
| | North America | 1247 | 0 |

[a] *The P values were obtained after a log-likelihood ratio test with 33 degrees of freedom*

***Table S4: Values of theta and the k parameter in the four genes within the 25kb chromosomal interval flanking the S-locus in 5'.***

| | **Genes** | **AT4G21410** | | **AT4G21400** | | **AT4G21390** | | **AT4G21380** | |
|---|---|---|---|---|---|---|---|---|---|
| **Species** | **Population** | **theta** | **k** | **theta** | **k** | **theta** | **k** | **theta** | **k** |
| *A. halleri* | Japan | 0.0049 | 0.31 | 0.0010 | 1.86 | 0.0058 | 2.35 | 0.0059 | 0.72 |
| | Nivelle | 0.0079 | 0.82 | 0.0023 | 2.2 | 0.0131 | 0.59 | 0.0062 | 1.86 |
| | Mortagne | 0.0071 | 7.93 | 0.0024 | 1.63 | 0.008 | 1.41 | 0.0067 | 4.32 |
| *A. lyrata* | Plech | 0.0108 | 3.35 | 0.0029 | 1.31 | 0.0123 | 1.99 | 0.0083 | 2.81 |
| | Spiterstulen | 0.0064 | 1.15 | 0.0013 | 2.01 | 0.0098 | 0.84 | 0.0066 | 0.63 |
| | North America | 0.0034 | 0.77 | 0.0014 | 2.54 | 0.0081 | 4.63 | 0.0089 | 1.75 |

*In this model, k measures the degree to which diversity is increased (k>1) or decreased (k<1) by the action of selection at each gene. The genetic diversity at each gene was estimated by theta.*

***Table S5: Values of theta and the k parameter in the seven genes within the 25kb chromosomal interval flanking the S-locus in 3'.***

| | **Genes** | **AT4G21350** | | **AT4G21340** | | **AT4G21330** | | **AT4G21323** | | **AT4G21310** | | **AT4G21300** | | **AT4G21280** | |
|---|---|---|---|---|---|---|---|---|---|---|---|---|---|---|---|
| **Species** | **Pop** | **theta** | **k** | **theta** | **k** | **theta** | **k** | **theta** | **k** | **theta** | **k** | **theta** | **k** | **theta** | **k** |
| *A. halleri* | Japan | 0.0089 | 0.46 | 0.0059 | 0.87 | 0.0063 | 2.09 | 0.0049 | 0.97 | 0.0104 | 1.46 | 0.0029 | 2.51 | 0.0059 | 0.03 |
| | Nivelle | 0.0102 | 2.21 | 0.0081 | 0.78 | 0.0048 | 4.5 | 0.0027 | 2.61 | 0.0114 | 2.46 | 0.0068 | 1.94 | 0.0109 | 0.16 |
| | Mortagne | 0.0133 | 1.21 | 0.007 | 0.98 | 0.0084 | 4.35 | 0.0037 | 2.68 | 0.0157 | 1.17 | 0.0065 | 3.5 | 0.0133 | 0.15 |
| *A. lyrata* | Plech | 0.015 | 2.28 | 0.0114 | 1.48 | 0.0074 | 1.32 | 0.0045 | 2.44 | 0.0125 | 1.08 | 0.0111 | 1.40 | 0.0144 | 0.61 |
| | Spiterstulen | 0.0064 | 2.02 | 0.0091 | 0.89 | 0.0067 | 0.49 | 0.0025 | 3.86 | 0.0092 | 1.53 | 0.0061 | 1.60 | 0.0132 | 0.53 |
| | North America | 0.0096 | 1.60 | 0.004 | 4.70 | 0.0081 | 1.42 | 0.0034 | 2.98 | 0.0111 | 4.33 | 0.0027 | 4.22 | 0.0049 | 0.87 |

*In this model, k measures the degree to which diversity is increased (k>1) or decreased (k<1) by the action of selection at each gene. The genetic diversity at each gene was estimated by theta.*

***Table S6: Variation of nucleotide polymorphism (π), minor allele frequency (MAF) and proportion of polymorphic sites across all sites in the S-flanking chromosomal intervals.***

| Species | Population | *S*-flanking regions | π | | MAF | | Proportion of polymorphic site | |
|---|---|---|---|---|---|---|---|---|
| | | | Value x10[3] [a] | Relative to controls [b] | Value [a] | Relative to controls [b] | Value x10[2] [a] | Relative to controls [b] |
| *A. halleri* | Japan | -75kb | 2.79 | 2.18 | 0.17 | 1.66 | 1.22 | 1.38 |
| | | -50kb | 2.36 | 1.84 | 0.07 | 0.68 | 2 | 2.26 |
| | | -25kb | **6.17** | 4.82 | 0.12 | 1.14 | **3.46** | 3.92 |
| | | 25kb | **7.41** | 5.78 | 0.17 | 1.59 | **3.2** | 3.62 |
| | | 50kb | 1.61 | 1.25 | 0.85 | 0.84 | 1.22 | 1.37 |
| | | 75kb | 3.35 | 2.61 | 0.2 | 1.97 | 1.3 | 1.46 |
| | Nivelle | -75kb | 5.51 | 1.35 | 0.22 | 0.98 | 1.9 | 1.46 |
| | | -50kb | 7.48 | 1.83 | 0.20 | 0.91 | 2.51 | 1.92 |
| | | -25kb | **14.14** | 3.46 | 0.21 | 0.94 | **4.8** | 3.67 |
| | | 25kb | **10.71** | 2.62 | 0.22 | 0.97 | **3.51** | 2.68 |
| | | 50kb | 4.71 | 1.15 | 0.20 | 0.92 | 1.62 | 1.24 |
| | | 75kb | 5.44 | 1.33 | 0.20 | 0.9 | 1.84 | 1.41 |
| | Mortagne | -75kb | 5.49 | 1.29 | 0.23 | 1.14 | 1.75 | 1.19 |
| | | -50kb | 7.58 | 1.79 | 0.24 | 1.16 | 2.36 | 1.61 |
| | | -25kb | **14.11** | 3.33 | 0.21 | 1.05 | **4.67** | 3.19 |
| | | 25kb | **10.21** | 2.41 | 0.19 | 0.92 | **3.75** | 2.56 |
| | | 50kb | 4.72 | 1.11 | 0.17 | 0.84 | 1.87 | 1.28 |
| | | 75kb | 4.17 | 0.98 | 0.14 | 0.69 | 2.03 | 1.39 |
| *A. lyrata* | Plech | -75kb | 8.97 | 1.18 | 0.15 | 0.89 | 3.94 | 1.32 |
| | | -50kb | 9.69 | 1.28 | 0.17 | 0.97 | 3.81 | 1.27 |
| | | -25kb | **14.1** | 1.86 | 0.16 | 0.93 | **5.9** | 1.97 |
| | | 25kb | **11.79** | 1.56 | 0.17 | 0.98 | **4.7** | 1.57 |
| | | 50kb | 10.69 | 1.41 | 0.17 | 0.99 | 4.15 | 1.39 |
| | | 75kb | 8.37 | 1.1 | 0.18 | 1.05 | 3.22 | 1.08 |
| | Spiterstulen | -75kb | 8.45 | 1.59 | 0.19 | 0.93 | 3.26 | 1.7 |
| | | -50kb | 5.83 | 1.09 | 0.19 | 0.93 | 2.17 | 1.14 |
| | | -25kb | **9.87** | 1.85 | 0.21 | 1.02 | **3.36** | 1.75 |
| | | 25kb | **9.98** | 1.87 | 0.16 | 0.76 | **4.41** | 2.3 |
| | | 50kb | 8.28 | 1.56 | 0.17 | 0.85 | **3.34** | 1.74 |
| | | 75kb | 6.17 | 1.16 | 0.19 | 0.94 | 2.29 | 1.19 |
| | North America | -75kb | 6.24 | 1.26 | 0.16 | 0.85 | 2.71 | 1.52 |
| | | -50kb | 6.36 | 1.28 | 0.21 | 1.10 | 2.16 | 1.21 |
| | | -25kb | **10.99** | 2.21 | 0.18 | 0.92 | **4.19** | 2.35 |
| | | 25kb | **10.09** | 2.03 | 0.23 | 1.21 | **3.28** | 1.84 |
| | | 50kb | 5.51 | 1.11 | 0.17 | 0.88 | 2.32 | 1.3 |
| | | 75kb | 5.25 | 1.06 | 0.21 | 1.12 | 1.73 | 0.97 |

[a] *Values departing from the 95% percentile of the distribution across control regions are shown in bold.* [b] *Ratio of the observed value in this chromosomal interval relative to the median of the 100 control regions.*

***Table S7: Linear model to evaluate the effect of distance to the S-locus (in kb) on nucleotide polymorphism (π) when measured on individual sites, either considering all sites in the flanking regions or 0-fold degenerate sites only.***

| | **π** | |
|---|---|---|
| | ***P* value** | **Linear effect** (per kb) |
| All sites | $<2x10^{-16}$ | $-8.65x10^{-5}$ |
| 0-fold degenerate sites | $2.75x10^{-15}$ | $-4.96x10^{-5}$ |

*Populations were included as random effects.*

***Table S8: Variation of nucleotide polymorphism (π), minor allele frequency (MAF) and proportion of polymorphic sites across 4-fold degenerate sites in the S-flanking chromosomal intervals.***

| **Species** | **Population** | ***S*-flanking regions** | **π** | | **MAF** | | **Proportion of polymorphic sites** | |
|---|---|---|---|---|---|---|---|---|
| | | | Value x10[3 a] | Relative to controls [b] | Value [a] | Relative to controls [b] | Value x10[2 a] | Relative to controls [b] |
| *A. halleri* | Japan | -75kb | 1.63 | 0.81 | 0.12 | 0.49 | 1.4 | 1.10 |
| | | -50kb | 4.48 | 2.22 | 0.06 | 0.45 | **4.34** | 3.42 |
| | | -25kb | 11.2 | 5.56 | 0.06 | 0.95 | **5.94** | 4.68 |
| | | 25kb | **15.9** | 7.85 | 0.15 | 1.22 | **6.95** | 5.48 |
| | | 50kb | 3.73 | 1.85 | 0.09 | 0.73 | 2.67 | 2.10 |
| | | 75kb | **14.9** | 7.40 | 0.22 | 1.77 | **4.97** | 3.92 |
| | Nivelle | -75kb | 7.42 | 1.11 | 0.17 | 0.76 | 2.8 | 1.29 |
| | | -50kb | 18.1 | 2.71 | 0.24 | 1.04 | 5.51 | 2.55 |
| | | -25kb | **26.9** | 4.03 | 0.19 | 0.83 | **9.84** | 4.55 |
| | | 25kb | **27.2** | 4.07 | 0.23 | 1.02 | **8.32** | 3.85 |
| | | 50kb | 10.4 | 1.55 | 0.26 | 1.16 | 2.91 | 1.35 |
| | | 75kb | 18.9 | 2.83 | 0.23 | 1.03 | 5.69 | 2.63 |
| | Mortagne | -75kb | 7.83 | 1.1 | **0.31** | **1.44** | 1.88 | 0.76 |
| | | -50kb | 17.6 | 2.47 | 0.27 | 1.24 | 4.94 | 1.99 |
| | | -25kb | **28.4** | 3.99 | 0.21 | 0.96 | **9.67** | 3.91 |
| | | 25kb | **23.7** | 3.33 | 0.19 | 0.90 | **8.49** | 3.43 |
| | | 50kb | 9.21 | 1.29 | 0.19 | 0.88 | 3.34 | 1.35 |
| | | 75kb | 14.2 | 2 | 0.16 | 0.76 | 5.87 | 2.37 |
| *A. lyrata* | Plech | -75kb | 1.49 | 0.96 | 0.16 | 0.84 | 6.16 | 1.07 |
| | | -50kb | 17 | 1.10 | 0.22 | 1.13 | 5.76 | 1 |
| | | -25kb | 30.6 | 1.98 | 0.19 | 0.98 | **11.3** | 1.96 |
| | | 25kb | **31.3** | 2.02 | 0.18 | 0.91 | **11.9** | 2.08 |
| | | 50kb | 17.2 | 1.11 | 0.16 | 0.83 | 6.84 | 1.19 |
| | | 75kb | 21.5 | 1.39 | 0.19 | 1.01 | 7.72 | 1.34 |
| | Spiterstulen | -75kb | 22.4 | 1.85 | 0.22 | 0.97 | **7.83** | 1.49 |
| | | -50kb | 18.9 | 1.57 | 0.20 | 0.93 | 6.23 | 1.71 |
| | | -25kb | 21.3 | 1.76 | 0.21 | 1.04 | 7.11 | 1.88 |
| | | 25kb | **26.3** | 2.17 | 0.18 | 0.86 | **10.1** | 2.43 |
| | | 50kb | 14.7 | 1.21 | 0.22 | 1.04 | 4.74 | 1.14 |
| | | 75kb | 17.7 | 1.46 | 0.20 | 0.93 | 6.12 | 1.47 |
| | North America | -75kb | 8.27 | 1 | 0.18 | 0.85 | 2.36 | 0.83 |
| | | -50kb | 3.88 | 0.47 | 0.26 | 1.24 | 1.17 | 0.41 |
| | | -25kb | **22** | 2.67 | 0.25 | 1.18 | **8.35** | 2.92 |
| | | 25kb | **23** | 2.79 | 0.25 | 1.18 | 6.90 | 2.41 |
| | | 50kb | 8.54 | 1.03 | 0.16 | 0.76 | 3.80 | 1.33 |
| | | 75kb | 18.3 | 2.21 | 0.30 | 1.39 | 4.62 | 1.61 |

[a] *Values departing from the 95% percentile of the distribution across control regions are shown in bold.* [b] *Ratio of the observed value in this chromosomal interval relative to the median of the 100 control regions.*

***Table S9: Variation of nucleotide polymorphism (π), minor allele frequency (MAF) and proportion of polymorphic sites across 0-fold degenerate sites in the S-flanking chromosomal intervals.***

| **Species** | **Population** | ***S*-flanking regions** | **π** | | **MAF** | | **Proportion of polymorphic site** | |
|---|---|---|---|---|---|---|---|---|
| | | | Value $x10^{3\ a}$ | Relative to controls [b] | Value [a] | Relative to controls [b] | Value $x10^{2\ a}$ | Relative to controls [b] |
| *A. halleri* | Japan | -75kb | 2.2 | 2.43 | 0.11 | 1.24 | 12.95 | 1.78 |
| | | -50kb | 1.25 | 1.39 | 0.05 | 0.51 | 13.99 | 1.93 |
| | | -25kb | 4.17 | 4.62 | 0.11 | 1.21 | **23.92** | 3.29 |
| | | 25kb | **5.95** | 6.59 | 0.16 | 1.69 | **26.6** | 3.66 |
| | | 50kb | 1.13 | 1.25 | 0.12 | 1.28 | 6.72 | 0.92 |
| | | 75kb | 3.17 | 3.51 | 0.17 | 1.78 | 14.63 | 2.01 |
| | Nivelle | -75kb | 3.03 | 1.09 | 0.18 | 0.84 | 11.95 | 1.29 |
| | | -50kb | 6.47 | 2.33 | 0.22 | 1.01 | 20.22 | 2.18 |
| | | -25kb | **8.02** | 2.89 | 0.18 | 0.84 | **30.88** | 3.33 |
| | | 25kb | **7.45** | 2.68 | 0.22 | 1.02 | **24.33** | 2.62 |
| | | 50kb | 3.3 | 1.19 | 0.21 | 0.99 | 10.74 | 1.16 |
| | | 75kb | 5.51 | 1.98 | 0.22 | 1.00 | 17.62 | 1.9 |
| | Mortagne | -75kb | 2.9 | 1.07 | 0.22 | 1.10 | 9.65 | 0.94 |
| | | -50kb | **7.16** | 2.64 | 0.26 | 1.29 | 21.35 | 2.07 |
| | | -25kb | **8.57** | 3.16 | 0.19 | 0.96 | **31.32** | 3.04 |
| | | 25kb | **6.89** | 2.54 | 0.19 | 0.95 | **25.67** | 2.49 |
| | | 50kb | 3.31 | 1.22 | 0.17 | 0.85 | 13.39 | 1.3 |
| | | 75kb | 3.23 | 1.19 | 0.11 | 0.56 | 18.77 | 1.82 |
| *A. lyrata* | Plech | -75kb | 3.28 | 0.57 | 0.16 | 0.95 | 14.86 | 0.65 |
| | | -50kb | **9.56** | 1.65 | 0.18 | 1.05 | **36.24** | 1.57 |
| | | -25kb | **10.69** | 1.84 | 0.14 | 0.83 | **44.16** | 1.92 |
| | | 25kb | 8.84 | 1.52 | 0.18 | 1.02 | 34.72 | 1.51 |
| | | 50kb | 7.1 | 1.22 | 0.16 | 0.95 | 29.29 | 1.27 |
| | | 75kb | 6.1 | 1.05 | 0.18 | 1.05 | 23.58 | 1.02 |
| | Spiterstulen | -75kb | 2.45 | 0.52 | 0.21 | 0.72 | 10.21 | 0.61 |
| | | -50kb | 6.97 | 1.48 | 0.15 | 0.68 | **30.5** | 1.83 |
| | | -25kb | 6.82 | 1.45 | 0.16 | 0.96 | 23.14 | 1.39 |
| | | 25kb | **8.87** | 1.88 | 0.18 | 0.82 | **34.74** | 2.08 |
| | | 50kb | **8.6** | 1.82 | 0.23 | 1.03 | **28.71** | 1.72 |
| | | 75kb | 4.18 | 0.89 | 0.17 | 0.76 | 16.97 | 1.02 |
| | North America | -75kb | 0.66 | 0.17 | 0.18 | 0.91 | 3.61 | 0.26 |
| | | -50kb | 5.23 | 1.37 | 0.20 | 1.03 | 18.97 | 1.38 |
| | | -25kb | **7.87** | 2.06 | 0.11 | 0.54 | **29.6** | 2.16 |
| | | 25kb | **8.49** | 2.22 | 0.21 | 1.07 | **28.89** | 2.11 |
| | | 50kb | 4.74 | 1.24 | 0.17 | 0.86 | 19.37 | 1.41 |
| | | 75kb | 4.98 | 1.3 | 0.31 | 1.57 | 12.63 | 0.92 |

[a] *Values departing from the 95% percentile of the distribution across control regions are shown in bold.* [b] *Ratio of the observed value in this chromosomal interval relative to the median of the 100 control regions.*

***Table S10: Identification of genes in S-flanking regions.***

| Gene name | *S*-flanking region | Gene ID NCBI | Symbol NCBI | Gene summary NCBI | Genomic coordinates [a] |
|---|---|---|---|---|---|
| AT4G21160 | +75kb | 827864 | ZAC | Calcium-dependent ARF-type GTPase activating protein family | Scaffold_7:9264247-9265403 |
| AT4G21170 | +75kb | 827865 | AT4G21170 | Tetratricopeptide repeat (TPR)-like superfamily protein | Scaffold_7:9268193-9268903 |
| AT4G21180 | +75kb | 827866 | ATERDJ2B | DnaJ / Sec63 Brl domains-containing protein | Scaffold_7:9273063-9275443 |
| AT4G21190 | +75kb | 827867 | emb1417 | Pentatricopeptide repeat (PPR) superfamily protein | Scaffold_7:9276639-9278563 |
| AT4G21192 | +75kb | 5008151 | AT4G21192 | Cytochrome c oxidase biogenesis protein Cmc1-like protein | Scaffold_7:9278695-9280631 |
| AT4G21200 | +75kb | 827868 | GA2OX8 | gibberellin 2-oxidase 8 | Scaffold_7:9281168-9283579 |
| AT4G21210 | +75kb | 827869 | RP1 | PPDK regulatory protein | Scaffold_7:9285282-9286317 |
| AT4G21215 | +50/75kb | 827870 | AT4G21215 | transmembrane protein | Scaffold_7:9299427-9300228 |
| AT4G21220 | +50Kb | 827871 | LpxD2 | Trimeric LpxA-like enzymes superfamily protein | Scaffold_7:9311284-9311889 |
| AT4G21230 | +50Kb | 827872 | CRK27 | cysteine-rich RLK (RECEPTOR-like protein kinase) 27 | Scaffold_7:9313820-9316515 |
| AT4G21240 | +50Kb | 827873 | AT4G21240 | F-box and associated interaction domains-containing protein | Scaffold_7:9317142-9329177 |
| AT4G21250 | +50Kb | 827874 | AT4G21250 | Sulfite exporter TauE/SafE family protein | Scaffold_7:9330684-9333743 |
| AT4G21270 | +50Kb | 827876 | ATK1 | kinesin 1 | Scaffold_7:9335860-9339460 |
| AT4G21280 | +25Kb/+50Kb | 827877 | PSBQA | photosystem II subunit QA | Scaffold_7:9376728-9377892 |
| AT4G21300 | +25Kb | 827878 | AT4G21300 | Tetratricopeptide repeat (TPR)-like superfamily protein | Scaffold_7:9380891-9382642 |
| AT4G21310 | +25Kb | 827879 | AT4G21310 | transmembrane protein, putative (DUF1218) | Scaffold_7:9384977-9385757 |
| AT4G21323 | +25Kb | 827881 | AT4G21323 | Subtilase family protein | Scaffold_7:9386059-9395172 |
| AT4G21330 | +25Kb | 827883 | DYT1 | basic helix-loop-helix (bHLH) DNA-binding superfamily protein | Scaffold_7:9395645-9396641 |
| AT4G21340 | +25Kb | 827884 | B70 | basic helix-loop-helix (bHLH) DNA-binding superfamily protein | Scaffold_7:9396651-9399212 |
| AT4G21350 | +25Kb | 827885 | PUB8 | plant U-box 8 | Scaffold_7:9400550-9402006 |
| AT4G21380 | -25Kb | 827890 | ARK3 | receptor kinase 3 | Scaffold_7:9402576-9406906 |
| AT4G21390 | -25Kb | 827891 | B120 | S-locus lectin protein kinase family protein | Scaffold_7:9407287-9409058 |
| AT4G21400 | -25Kb | 827892 | CRK28 | cysteine-rich RLK (RECEPTOR-like protein kinase) 28 | Scaffold_7:9413560-9414613 |
| AT4G21410 | -25Kb/-50Kb | 827893 | CRK29 | cysteine-rich RLK (RECEPTOR-like protein kinase) 29 | Scaffold_7:9414947-9417503 |
| AT4G21430 | -50Kb | 827895 | B160 | protein B160 | Scaffold_7:9417579-9419697 |
| AT4G21440 | -50Kb | 826916 | MYB102 | MYB-like 102 | Scaffold_7:9422970-9426554 |
| AT4G21445 | -50Kb | 825893 | AT4G21445 | receptor-interacting protein | Scaffold_7:9428598-9430280 |
| AT4G21450 | -75Kb | 826300 | AT4G21450 | PapD-like superfamily protein | Scaffold_7:9430712-9434329 |
| AT4G21470 | -75Kb | 828232 | FMN/FHY | riboflavin kinase/FMN hydrolase | Scaffold_7:9435110-9442269 |
| AT4G21480 | -75Kb | 828233 | STP12 | sugar transporter protein 12 | Scaffold_7:9444465-9445786 |
| AT4G21490 | -75Kb | 828234 | NDB3 | NAD(P)H dehydrogenase B3 | Scaffold_7:9445909-9449298 |
| AT4G21500 | -75Kb | 828235 | AT4G21500 | transmembrane protein | Scaffold_7:9449750-9451507 |
| AT4G21520 | -75Kb | 828237 | AT4G21520 | Transducin/WD40 repeat-like superfamily protein | Scaffold_7:9451640-9454210 |

[a] *The genomic coordinates on scaffold 7 of A. lyrata.*

***Table S11: Phenotypes associated with genes in S-flanking regions of 25kb presenting an increase of polymorphism in different studies.***

| **Gene name** | **Species** | **Reference study** | **Phenotype associated** |
|---|---|---|---|
| AT4G21300 | *Citrus sinensis* | *Lavagi-Craddock et al. 2022* | Response to viroid |
| AT4G21330 | *Arabidopsis thaliana* | *Zhang et al. 2006* | Anther morphology |
| AT4G21330 | *C. sinensis* | *Ma et al. 2016* | Magnesium-deficiency in leaves |
| AT4G21330 | *Gossypium hirsutum* | *Hamid et al. 2019* | Pollen sterility |
| AT4G21380 | *A. thaliana* | *Dwyer et al. 1994* | Root development |
| AT4G21380 | *A. thaliana* | *Pastuglia et al. 2002* | Plant defence response |
| AT4G21380 | *A. thaliana* | *Mondal et al. 2021* | Abiotic stress response |
| AT4G21390 | *A. thaliana* | *Chae et al. 2009* | Abiotic stress response |
| AT4G21390 | *A. thaliana* | *Meier et al. 2010* | Response to pathogens |
| AT4G21390 | *A. thaliana* | *Kissen et al. 2016* | Heat stress, oxidative stress, plant defence responses |
| AT4G21390 | *A. thaliana* | *Noman et al. 2019* | Drought tolerance |
| AT4G21390 | *A. thaliana* | *Mondal et al. 2021* | Abiotic stress responses |
| AT4G21390 | *Zea mays* | *Welgemoed et al. 2020* | Resistance to Gray Leaf Spot disease |
| AT4G21390 | *Z. mais* | *Aglyamova et al. 2020* | Root development |
| AT4G21390 | *Arachis stenosperma* | *Martins et al. 2022* | Responses to UV-C exposure |
| AT4G21390 | *Lolium perenne* | *Zhang et al, 2022* | Plant defence response |
| AT4G21400 | *A. thaliana* | *Engelmann et al. 2008* | Response to pathogens |
| AT4G21400 | *A. thaliana* | *Yadeta et al. 2017* | Immunity |
| AT4G21400 | *A. thaliana* | *Lauss et al. 2018* | Hybrid heterosis |
| AT4G21410 | *A. thaliana* | *Engelmann et al. 2008* | Response to pathogens |
| AT4G21410 | *A. thaliana* | *Yadeta et al. 2017* | Immunity |
| AT4G21410 | *A. thaliana* | *Lauss et al. 2018* | Hybrid heterosis |

***Table S12: Genomic location and size of the one hundred 25kb control regions defined in the* A. halleri *genome and their corresponding location in the* A. lyrata *genome, as determined by YASS.***

| Control region in *A. halleri* genome | | | Corresponding region studied in *A. lyrata* genome | | | |
|---|---|---|---|---|---|---|
| **Chromosome** | **First position** | **Last position** | **Chromosome** | **First position** | **Last position** | **Size (bp)** |
| scaffold1 | 714746 | 739746 | 5 | 19398229 | 19412882 | 14653 |
| scaffold10 | 375896 | 400896 | 3 | 7536232 | 7560491 | 24259 |
| scaffold106 | 225230 | 250230 | 4 | 16834544 | 16844691 | 10147 |
| scaffold11 | 386718 | 411718 | 8 | 19413760 | 19431480 | 17720 |
| scaffold116 | 198360 | 223360 | 7 | 727463 | 751667 | 24204 |
| scaffold119 | 166294 | 191294 | 8 | 21802390 | 21825492 | 23102 |
| scaffold120 | 88820 | 113820 | 1 | 32869077 | 32875910 | 6833 |
| scaffold121 | 99101 | 124101 | 6 | 6851532 | 6867611 | 16079 |
| scaffold128 | 11070 | 36070 | 7 | 9110002 | 9121284 | 11282 |
| scaffold129 | 115566 | 140566 | 4 | 22523499 | 22539888 | 16389 |
| scaffold136 | 65513 | 90513 | 3 | 9589443 | 9614893 | 25450 |
| scaffold137 | 69718 | 94718 | 5 | 16514721 | 16532625 | 17904 |
| scaffold138 | 211414 | 236414 | 3 | 3614206 | 3638620 | 24414 |
| scaffold142 | 5686 | 30686 | 7 | 21879403 | 21897701 | 18298 |
| scaffold145 | 190577 | 215577 | 7 | 1531420 | 1570539 | 39119 |
| scaffold150 | 202351 | 227351 | 5 | 12253381 | 12265905 | 12524 |
| scaffold152 | 184519 | 209519 | 8 | 16926540 | 16961242 | 34702 |
| scaffold155 | 166053 | 191053 | 3 | 3261529 | 3277110 | 15581 |
| scaffold162 | 8620 | 33620 | 7 | 9568346 | 9607514 | 39168 |
| scaffold166 | 16582 | 41582 | 2 | 3244155 | 3269796 | 25641 |
| scaffold170 | 75064 | 100064 | 7 | 12975344 | 12992621 | 17277 |
| scaffold171 | 218981 | 243981 | 1 | 11635934 | 11651879 | 15945 |
| scaffold173 | 151316 | 176316 | 5 | 4003159 | 4032530 | 29371 |
| scaffold174 | 176210 | 201210 | 6 | 4249736 | 4277114 | 27378 |
| scaffold177 | 211995 | 236995 | 3 | 12180930 | 12186261 | 5331 |
| scaffold179 | 152543 | 177543 | 7 | 23594987 | 23610811 | 15824 |
| scaffold18 | 604881 | 629881 | 4 | 20412360 | 20431225 | 18865 |
| scaffold181 | 155971 | 180971 | 6 | 23226457 | 23244394 | 17937 |
| scaffold194 | 3536 | 28536 | 8 | 1401121 | 1426726 | 25605 |
| scaffold200 | 125357 | 150357 | 1 | 83447 | 120180 | 36733 |
| scaffold201 | 109767 | 134767 | 4 | 21525149 | 21561242 | 36093 |

| | | | | | | |
|---|---|---|---|---|---|---|
| scaffold207 | 22822 | 47822 | 8 | 1193675 | 1218564 | 24889 |
| scaffold21 | 443229 | 468229 | 6 | 24844808 | 24897223 | 52415 |
| scaffold218 | 108209 | 133209 | 3 | 21611295 | 21635129 | 23834 |
| scaffold224 | 28462 | 53462 | 2 | 12848081 | 12862599 | 14518 |
| scaffold225 | 117783 | 142783 | 3 | 22174587 | 22186666 | 12079 |
| scaffold227 | 208256 | 233256 | 8 | 13458265 | 13481329 | 23064 |
| scaffold229 | 56619 | 81619 | 1 | 23286221 | 23293991 | 7770 |
| scaffold23 | 327652 | 352652 | 2 | 16035207 | 16049270 | 14063 |
| scaffold232 | 119994 | 144994 | 7 | 433865 | 457338 | 23473 |
| scaffold235 | 5432 | 30432 | 7 | 23331768 | 23363142 | 31374 |
| scaffold239 | 50880 | 75880 | 6 | 10797019 | 10810414 | 13395 |
| scaffold242 | 59705 | 84705 | 1 | 1418754 | 1430112 | 11358 |
| scaffold250 | 128700 | 153700 | 8 | 17573303 | 17600436 | 27133 |
| scaffold26 | 414726 | 439726 | 7 | 3634264 | 3651699 | 17435 |
| scaffold27 | 415774 | 440774 | 7 | 11175491 | 11196741 | 21250 |
| scaffold273 | 72568 | 97568 | 5 | 1653956 | 1664424 | 10468 |
| scaffold28 | 423509 | 448509 | 7 | 7622615 | 7639343 | 16728 |
| scaffold281 | 45642 | 70642 | 1 | 15688656 | 15731735 | 43079 |
| scaffold29 | 88776 | 113776 | 2 | 17654931 | 17671824 | 16893 |
| scaffold3 | 341239 | 366239 | 2 | 14262838 | 14285032 | 22194 |
| scaffold314 | 91953 | 116953 | 1 | 12266331 | 12285752 | 19421 |
| scaffold316 | 12224 | 37224 | 7 | 9705209 | 9725734 | 20525 |
| scaffold317 | 16150 | 41150 | 5 | 17816355 | 17837342 | 20987 |
| scaffold32 | 546985 | 571985 | 3 | 8899593 | 8920135 | 20542 |
| scaffold323 | 72038 | 97038 | 7 | 24099129 | 24102364 | 3235 |
| scaffold33 | 17665 | 42665 | 6 | 437309 | 469105 | 31796 |
| scaffold330 | 61130 | 86130 | 8 | 22307313 | 22313905 | 6592 |
| scaffold34 | 813865 | 838865 | 4 | 15130559 | 15152366 | 21807 |
| scaffold358 | 32810 | 57810 | 3 | 11023648 | 11050966 | 27318 |
| scaffold36 | 635729 | 660729 | 8 | 22233394 | 22257401 | 24007 |
| scaffold361 | 190619 | 215619 | 4 | 4489852 | 4494894 | 5042 |
| scaffold37 | 226418 | 251418 | 5 | 15605860 | 15630386 | 24526 |
| scaffold4 | 82927 | 107927 | 6 | 10346324 | 10381808 | 35484 |
| scaffold40 | 186691 | 211691 | 4 | 19035410 | 19056081 | 20671 |
| scaffold42 | 530313 | 555313 | 1 | 4448641 | 4467582 | 18941 |
| scaffold439 | 79475 | 104475 | 4 | 4120140 | 4152041 | 31901 |

| | | | | | | |
|---|---|---|---|---|---|---|
| scaffold44 | 20781 | 45781 | 4 | 11117870 | 11132007 | 14137 |
| scaffold444 | 4652 | 29652 | 6 | 21259002 | 21291721 | 32719 |
| scaffold451 | 217461 | 242461 | 7 | 6691332 | 6704495 | 13163 |
| scaffold46 | 101487 | 126487 | 3 | 4048724 | 4060778 | 12054 |
| scaffold47 | 44051 | 69051 | 3 | 2834327 | 2843749 | 9422 |
| scaffold48 | 357354 | 382354 | 5 | 951841 | 975828 | 23987 |
| scaffold485 | 56582 | 81582 | 8 | 2782403 | 2805264 | 22861 |
| scaffold49 | 428306 | 453306 | 7 | 6037403 | 6066170 | 28767 |
| scaffold505 | 31658 | 56658 | 3 | 9289036 | 9316360 | 27324 |
| scaffold51 | 393946 | 418946 | 4 | 22852372 | 22871075 | 18703 |
| scaffold518 | 67369 | 92369 | scaffold_41 | 2689 | 11067 | 8378 |
| scaffold52 | 332919 | 357919 | 5 | 15029455 | 15051052 | 21597 |
| scaffold54 | 221637 | 246637 | 3 | 939725 | 967894 | 28169 |
| scaffold553 | 11337 | 36337 | 2 | 3555776 | 3569114 | 13338 |
| scaffold573 | 37459 | 62459 | 7 | 23035137 | 23062551 | 27414 |
| scaffold599 | 19858 | 44858 | 7 | 19598096 | 19603720 | 5624 |
| scaffold6 | 832830 | 857830 | 1 | 5098531 | 5118542 | 20011 |
| scaffold60 | 31634 | 56634 | 3 | 7830576 | 7839059 | 8483 |
| scaffold612 | 8199 | 33199 | 6 | 10307793 | 10331203 | 23410 |
| scaffold64 | 278151 | 303151 | 6 | 2224735 | 2252345 | 27610 |
| scaffold65 | 758674 | 783674 | 1 | 8784710 | 8818666 | 33956 |
| scaffold653 | 40454 | 65454 | 8 | 13584566 | 13621779 | 37213 |
| scaffold66 | 290017 | 315017 | 6 | 1147402 | 1165983 | 18581 |
| scaffold67 | 295039 | 320039 | 6 | 7343155 | 7365804 | 22649 |
| scaffold71 | 293597 | 318597 | 1 | 1069475 | 1082780 | 13305 |
| scaffold76 | 328971 | 353971 | 6 | 4138280 | 4163654 | 25374 |
| scaffold79 | 484435 | 509435 | 8 | 639549 | 668410 | 28861 |
| scaffold8 | 358549 | 383549 | 8 | 21436030 | 21456224 | 20194 |
| scaffold81 | 121244 | 146244 | 1 | 9779214 | 9802443 | 23229 |
| scaffold84 | 274392 | 299392 | 3 | 1082652 | 1101060 | 18408 |
| scaffold86 | 186170 | 211170 | 3 | 4340346 | 4364458 | 24112 |
| scaffold9 | 208753 | 233753 | 6 | 8241007 | 8260480 | 19473 |
| scaffold93 | 271013 | 296013 | 7 | 4576624 | 4588042 | 11418 |

Figures

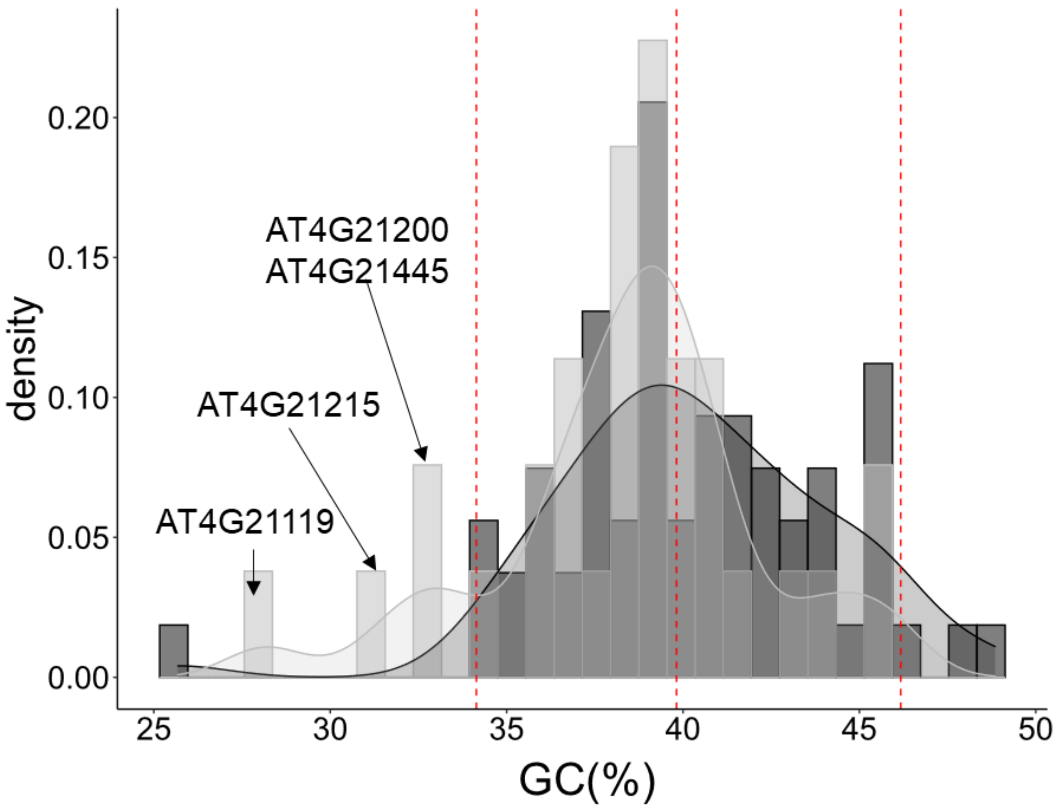


***Figure S1: Comparaison of the GC content of genes in control vs. S-flanking regions in A. lyrata.***

*Distribution curve estimated by ggplot package in R with default parameters. The red dashed lines represent the values in 2.5, 50 and 97.5% of the GC contained in control genes.*

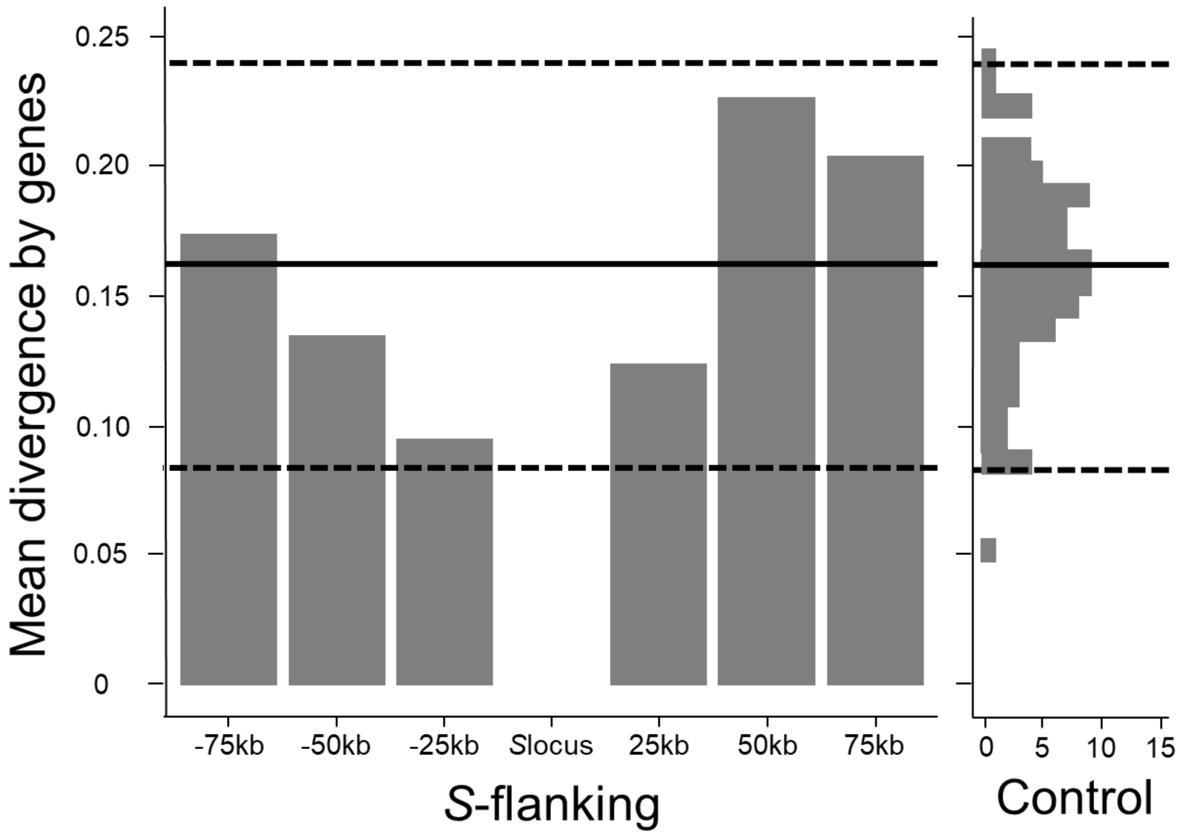


***Figure S2 : Mean A. thaliana - A. lyrata divergence in genes distributed around the S-locus and across the control regions from throughout the genome.*** *Bars represent the mean value of divergence (proportion of divergent sites in genes between the two species) obtained in each region of 25kb around the S-locus. The distribution of the mean divergence in genes in the 100 control regions is represented by the histogram on the right. The median value of the distribution in control regions is represented by the black line and the 97.5% and 2.5% interval by the dashed lines.*

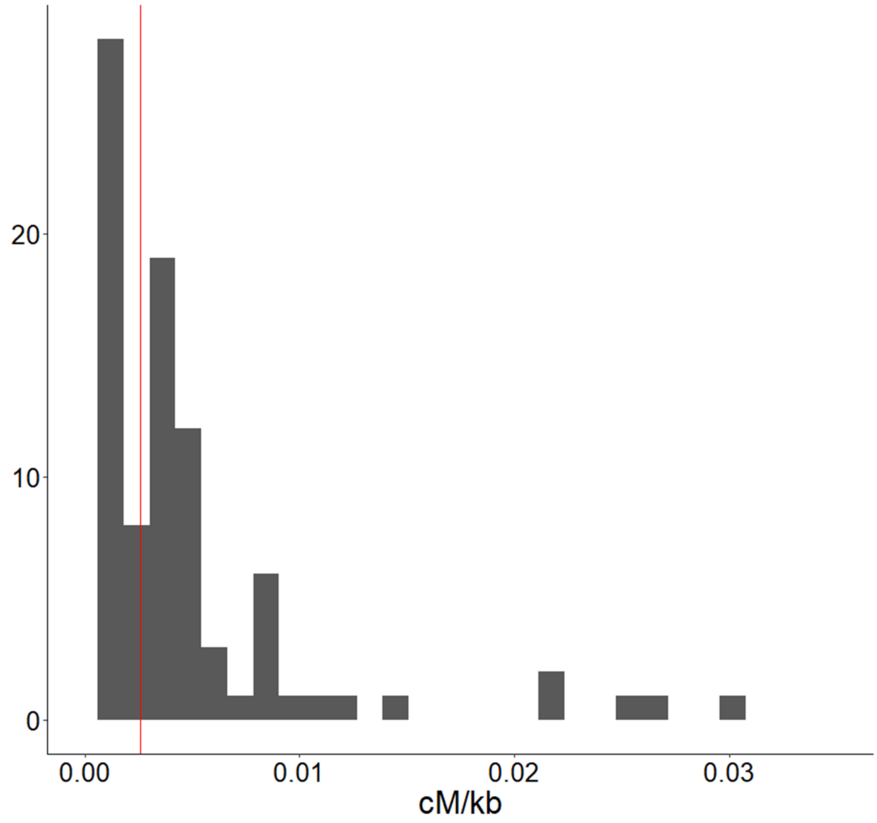


***Figure S3: Comparison of the recombination rates (cM/kb) in A. lyrata in genomic intervals containing the control regions (grey bars) and the S-locus (red line).***

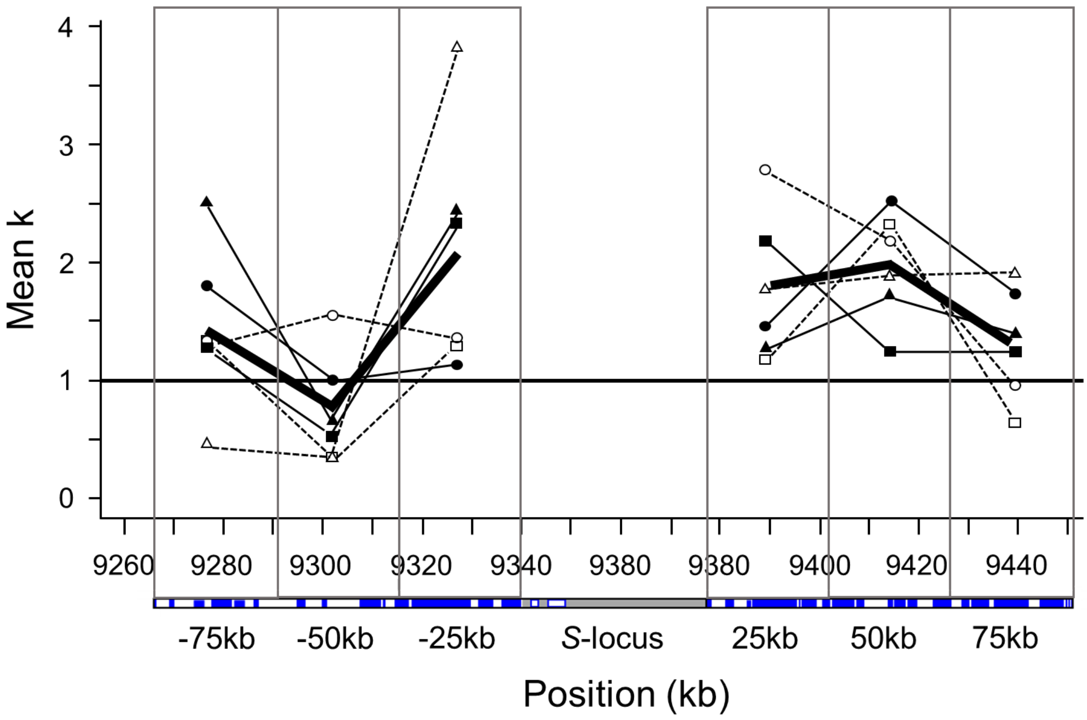


***Figure S4 : Mean selection parameter (k) estimated by MLHKA for genes along the S-flanking regions.*** *The thick solid line represents the mean value of k obtained across the six sample sets. The thin solid lines represent the A. lyrata sample sets (square=Plech, circle=Spiterstulen, triangle=North America). The dashed lines represent the A. halleri sample sets (open square=Japan, open circle=Nivelle, open triangle=Mortagne). The threshold value of k=1 (no selection) is represented by the horizontal black line. The graduation along the X axis represents nucleotide positions along chromosome 7 of the A. lyrata genome assembly (in kb).*

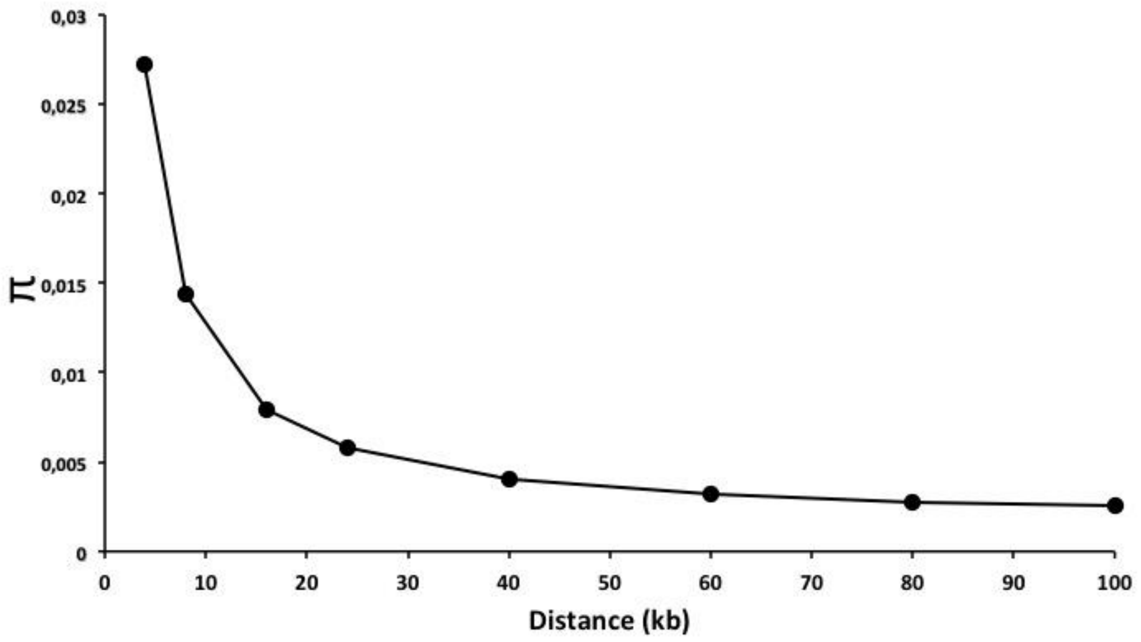


***Fig. S5 : Expected neutral nucleotide diversity in a region partially linked to a locus under strong balancing selection as a function of distance from the selected locus, according to Takahata & Satta (1998)'s equations.*** *Balancing selection is overdominance with s=1, Ne=2500, u = rate of introduction of new selected alleles = 1.10-9. The mutation rate at the neutral locus is v = 1.5 10-7. The local recombination rate is 2.5 cM/Mb.*

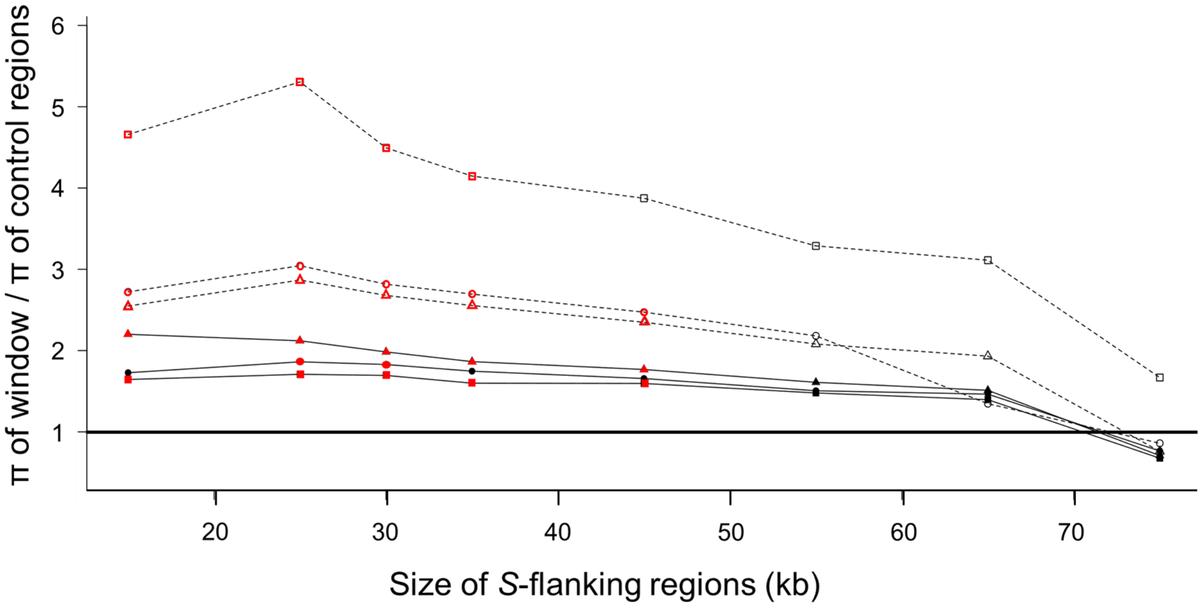


***Figure S6 : The elevation of nucleotide polymorphism (π) in the S-flanking regions as compared to the control regions is sensitive to the choice of the window size.*** *The windows with a π higher than the 95% percentiles of the control region are represented by red symbols. The solid lines represent the A. lyrata sample sets (square=Plech, circle=Spiterstulen, triangle=North America). The dashed lines represent the A. halleri sample sets (open square=Japan, open circle=Nivelle, open triangle=Mortagne). The threshold value of 1 (same π in S-linked vs control regions) is represented by the horizontal black line.*

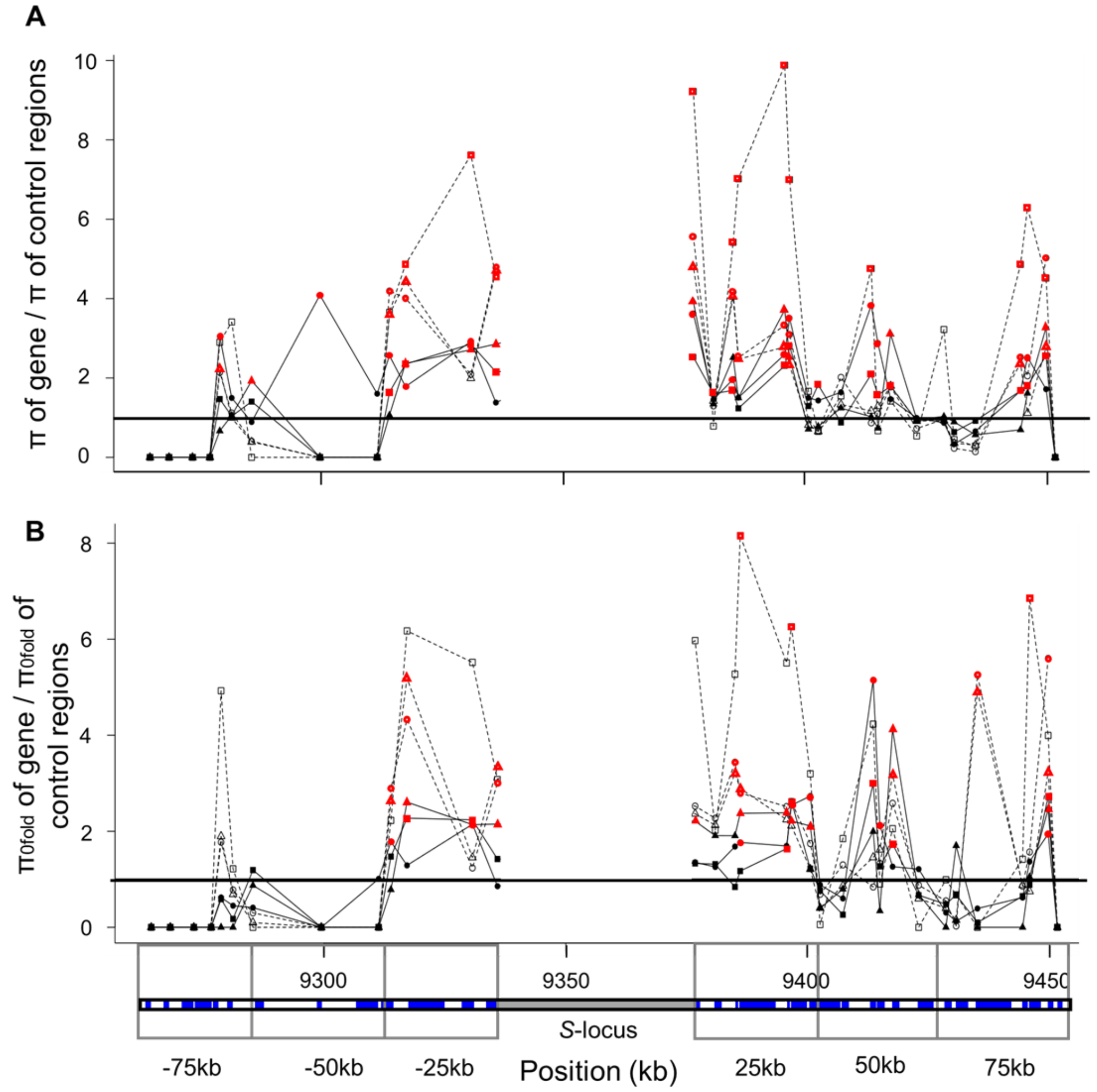


***Figure S7 : Nucleotide polymorphism (π) of genes in the S-flanking regions relative to the median π in the control regions.*** *A) Total polymorphism (all sites included). B) 0-fold degenerate sites only. The solid lines represent the A. lyrata sample sets (square=Plech, circle=Spiterstulen, triangle=North America). The dashed lines represent the A. halleri sample sets (open square=Japan, open circle=Nivelle, open triangle=Mortagne). The threshold value of 1 (same π in S-linked vs control regions) is represented by the horizontal black line. The genes whose π value is higher than the 95% percentile of the control regions are represented by red symbols.*

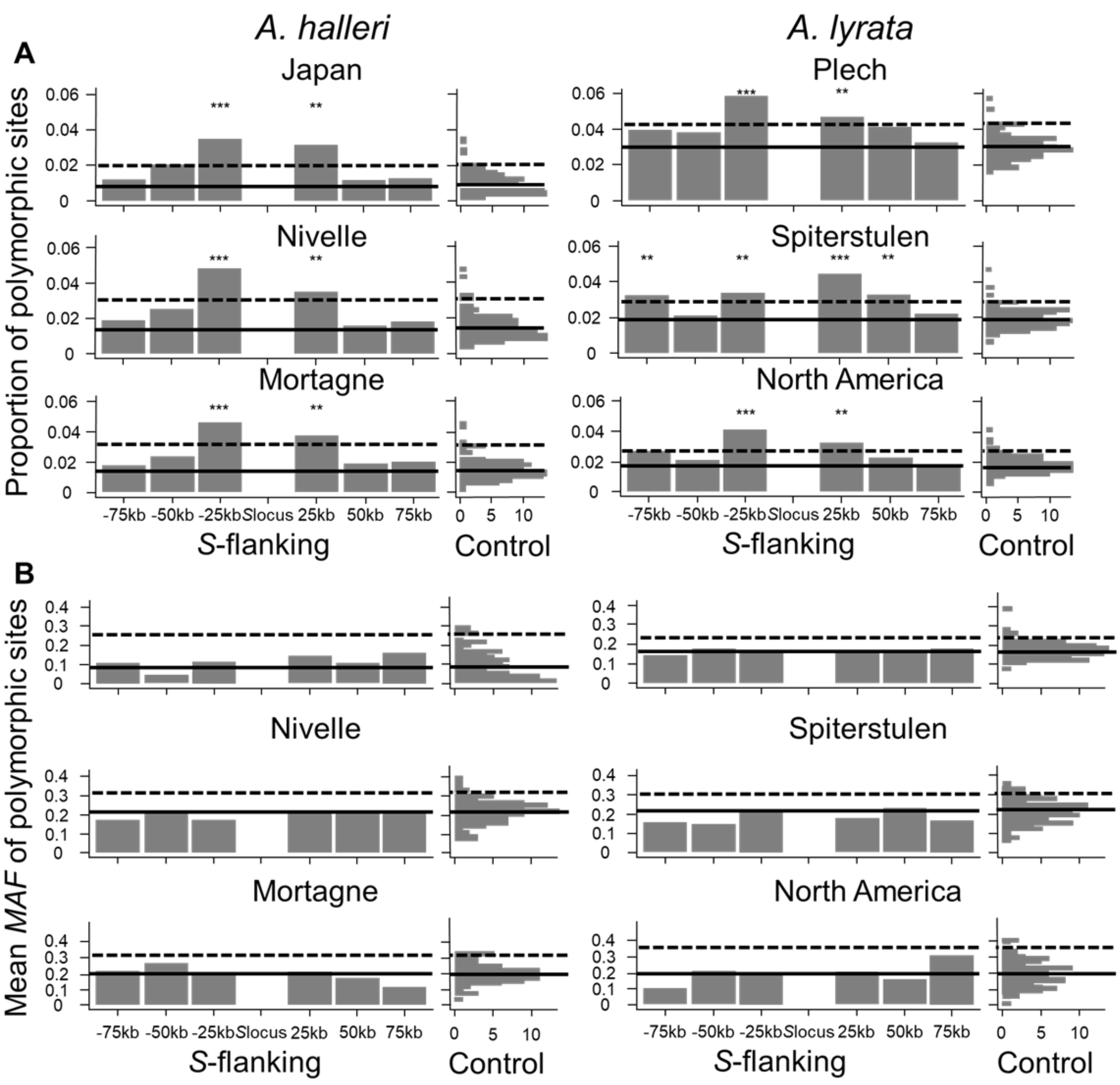


***Figure S8: Pattern of polymorphism in sites around the S-locus and across the control regions from throughout the genome.*** *A) Proportion of polymorphic sites. B) Mean MAF in polymorphic sites. Bars represent the proportion of polymorphic sites (A) or the mean MAF (B) in non-overlapping regions of 25kb around the S-locus. The distributions (count) of the same respective statistic in the 100 control regions are represented by a vertical histogram on the right. The 95% percentile of the distributions is represented by dashed lines. The median value of the distribution in control regions is represented by black lines. *** = observed value above the 99% percentile of control regions, ** = observed value above the 97,5% percentile of control regions, * = observed value above the 95% percentile of control regions.*
